# Bacteriophage uptake by Eukaryotic cell layers represents a major sink for phages during therapy

**DOI:** 10.1101/2020.09.07.286716

**Authors:** Marion C. Bichet, Wai Hoe Chin, William Richards, Yu-Wei Lin, Laura Avellaneda-Franco, Catherine A. Hernandez, Arianna Oddo, Oleksandr Chernyavskiy, Volker Hilsenstein, Adrian Neild, Jian Li, Nicolas Hans Voelcker, Ruzeen Patwa, Jeremy J. Barr

**Affiliations:** School of Biological Sciences, Monash University, Clayton Campus, VIC, 3800, Australia; Biomedicine Discovery Institute and Department of Microbiology, Monash University, Clayton, VIC, Australia; Department of Integrative Biology, University of California, Berkeley, Berkeley, CA, USA; Faculty of Pharmacy and Pharmaceutical Sciences, Monash University, Parkville Campus, VIC, 3800, Australia; Monash Micro Imaging, Monash University, Clayton Campus, VIC, 3800, Australia; Department of Mechanical and Aerospace Engineering, Monash University, Clayton Campus, VIC, 3800, Australia

## Abstract

For over 100 years, bacteriophages have been known as viruses that infect bacteria. Yet it is becoming increasingly apparent that bacteriophages, or phages for short, have tropisms outside their bacterial hosts. During phage therapy, high doses of phages are directly administered and disseminated throughout the body, facilitating broad interactions with eukaryotic cells. Using live cell imaging across a range of cell lines we demonstrate that cell type plays a major role in phage internalisation and that smaller phages (< 100 nm) are internalised at higher rates. Uptake rates were validated under physiological shear stress conditions using a microfluidic device that mimics the shear stress to which endothelial cells are exposed to in the human body. Phages were found to rapidly adhere to eukaryotic cell layers, with adherent phages being subsequently internalised by macropinocytosis and functional phages accumulating and stably persisting intracellularly. Finally, we incorporate these results into an established pharmacokinetic model demonstrating the potential impact of phage accumulation by these cell layers, which represents a major sink for circulating phages in the body. Understanding these interactions will have important implications on innate immune responses, phage pharmacokinetics, and the efficacy of phage therapy.

## Introduction

Phages, short for bacteriophages, are viruses that infect bacteria and are the most abundant life form on the planet (1–3). Phages are found ubiquitously in the environment and are a major contributor to global genetic diversity (4–6). Our bodies harbour a large number of phages, and, together with their bacterial hosts, they constitute a key component of our gut microbiome (7). The gut carries the largest aggregation of phages in the body, with an estimated 2 × 10^12^ phages present in the average human colon (4,8,9). These phages are constantly interacting with gut bacteria, as well as the epithelial cell layers of the gut (10). Phages are detected in the circulatory systems of the body, suggesting they are capable of translocating from the gut and penetrating throughout the body (8,11). Once past the gut barrier phages are able to penetrate cell layers and major organs of the body; being found in classically sterile regions such as the blood, serum, organs and even the brain (8,10–19). Numerous mechanisms pertaining to the transport of phages across epithelial barriers have been proposed (11,16), including the ‘leaky gut’ where phages bypass cell barriers at sites of damage and inflammation (20,21), and receptor-mediated endocytosis (22,23). Recently, a non-specific mechanism for phage uptake and transport across epithelial cell layers was proposed by Nguyen and colleagues, whereby epithelial cells uptake phages via macropinocytosis and preferentially transcytose phages from the apical surface toward the basolateral side of the cell (8,24). Macropinocytosis is a broad mechanism describing the enclosure of media within ruffles in cells’ membrane, prior to internalising the media, and any phages it may contain, within the cells. Despite their prevalence in the human body, phage’s capacity to interact with and influence eukaryotic cells remains largely unknown. These interactions can have important implications during phage therapy.

Phage therapy is a promising alternative to treat pathogenic bacterial infections. In Eastern Europe, phage therapy was widely used since its discovery in 1917 (1), whereas in Western countries, phage therapy was largely abandoned in favour of antibiotics (25–28). However, with the rise of antimicrobial resistance as one of the greatest threats to human health, phage therapy is being re-established as a potential treatment option for difficult-to-treat, antibiotic resistant, bacterial infections (25). In order for phage therapy to be effective, phages must first be administered to the site of infection. Administration routes include, intravenous (IV) or intraperitoneal (IP) to treat septicaemia; orally to treat gastrointestinal infections; intranasal or inhalation to treat respiratory infections; or topically for cutaneous infections (29,30). The administration route and bioavailability of phages needs to be carefully taken into account in order to achieve favourable efficacy *in vivo*.

In contrast to conventional drugs, phages are unique therapeutic agents capable of self-replicating and maintaining titres in the body (30–33). As such, there is a lack of knowledge regarding phage pharmacokinetics and pharmacodynamics (31). Following administration, two major pharmacokinetic factors important for the efficacy of phage therapy are accessibility and clearance. First, natural barriers such as cell layers and mucus can decrease accessibility of phages to sites of infection, thereby necessitating the administration of higher doses to achieve a favourable therapeutic effect. Second, phage clearance has been reported to occur rapidly; sometimes within just minutes to hours following parenteral administration in animal models and patients (14,34–40). Phage clearance within the body is thought to be mediated by three main components: 1) Phagocytic cells (41), 2) The Mononuclear Phagocytic System (MPS; which was also previously called the reticuloendothelial system, or RES) which includes the liver and spleen that filter out and remove phages from circulation (42), and 3) Phage neutralizing antibodies, although it is still unclear how effective and rapidly produced these anti-phage antibodies are (31,43). Due to these complications, it is difficult to predict how phages will behave in the body when administered and ultimately whether phage therapy will be successful.

One underexplored aspect of phage therapy is the non-specific interactions between phages and the cell layers of the body. During therapy, large quantities of phage monocultures or cocktails are administered to patients in order to maintain a killing titre to combat a bacterial infection. Once within the body, these phages can have very short half-lives and are actively removed or inactivated by the body (38–40,44). Following administration during phage therapy, epithelial and endothelial cell layers are amongst the first and most abundant phage-eukaryote interactions. Here, we present new insights into phage-mammalian cell adherence, uptake, and trafficking, via *in vitro* tissue culture cell layers. Our results suggest that cell layers of the human body represent a major and unaccounted sink for exogenously administered phages. Put within the context of phage therapy, the interaction between eukaryotic cells and exogenous phages have important implications for phage administration, dosing, and pharmacokinetics.

## Results

### The rate of phage uptake varies depending on the cell type

To better understand phage-eukaryote interactions, we used seven *in vitro* tissue culture cell lines that were selected to be broadly representative of different tissues types within the body, and examined their interactions with ultrapure high titre monocultures of T4 phage. Using real time live-cell imaging on a confocal microscope with a sensitive hybrid detector (HyD), we visualised the interaction and subsequent internalisation of phage particles within eukaryotic cells. Cells were first grown on glass bottom slides for two days to generate an ∼ 80% confluent cell layer, followed by fluorescence staining of the nucleus and plasma membrane. T4 phages were prepared using the Phage-on-Tap method (45), labelled using SYBR-Gold, subsequently washed to remove residual stain, and then directly applied to cell layers observed using live cell imaging.

Phages were visualised in real time being engulfed and trafficked through all seven of the different cell lines over a two-hour period (Fig. 1a and Supplemental movies SM1-7). The cell types tested include: epithelial cells – HeLa, A549 and HT29, from the cervix, lung and colon, respectively; fibroblast cells – MDCK-I and BJ, from dog kidney and human skin, respectively; the endothelial cell line – HUVEC from umbilical vein; and monocyte-induced macrophages – THP-1 cells (Fig. 1b). The increase in green fluorescence over time corresponds to the uptake and accumulation of fluorescently labelled T4 phages by the cells. We saw the first evidence of phage accumulation within cells occurring around 30 minutes, with continued accumulation over the following 90 minutes.

**Figure 1.**
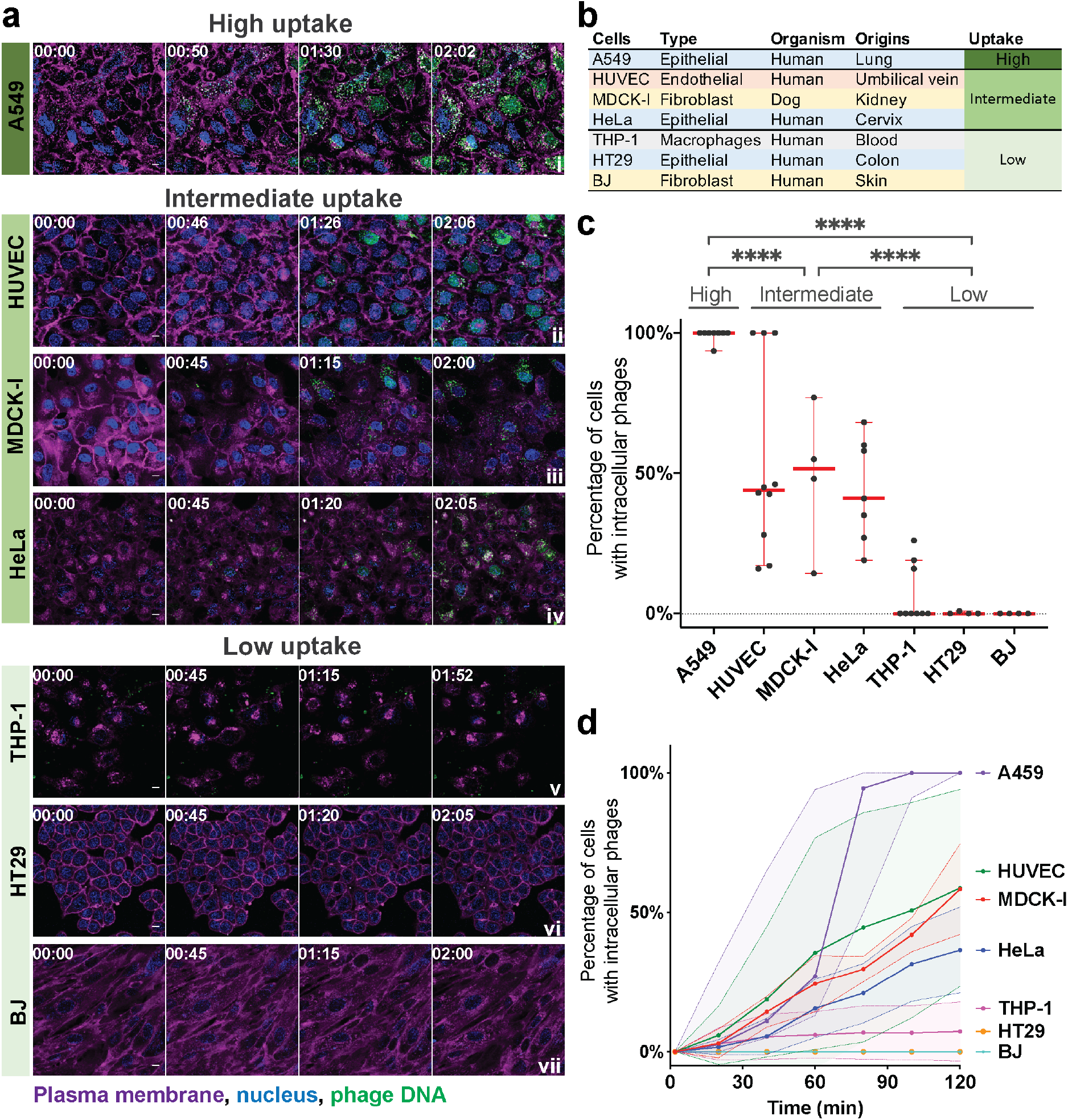
Uptake and internalisation of phages varies across cell type. (**a**) T4 phage was fluorescently labelled, applied to different cell lines and incubated for two hours on a glass bottom slide. Cells were stained with Hoechst 33342 nucleus stain (blue), CellMask plasma membrane stain (magenta) and T4 phages labelled with SYBR-Gold DNA-complexing stain (green). Using real-time microscopy, one image was acquired every two minutes. Scale bar: 10 µm; Timing: hours:minutes. (**b**) Table of cell lines used in this study, their cell type, organism, organ origins and category of uptake. Cells lines are ranked as high, intermediate, and low uptake. (**c**) Percentage of cells containing intracellular phages at the two-hour time point. Scatter plots show medians of percentage of cells with intracellular phages; error bars represent 95% confidence intervals; each dot represents one Field of View (FOV). Cells lines are ranked as high, intermediate and low uptake. P-values between the different groups calculated from a one-way ANOVA, shown as stars (F (2, 43) = 71.23; P < 0.0001: ****). (**d**) Percentage of cells containing phages represented across time. For each video the number of cells with and without intracellular phages in a FOV were manually counted every ten minutes (A549 *n* = 6; HUVEC *n* = 8; MDCK-I *n* = 3; HeLa *n* = 6; THP-1 *n* = 8; HT29 *n* = 3; BJ *n* = 3). Curve plots show medians of percentage of cells with intracellular phages; error bars represent 95% confidence intervals in the shaded area for each curve; each point represents one video over time.

We observed large variation in the uptake of phages across the seven cell types investigated. To quantify this, cells containing intracellular phages were manually counted and compared with the total number of cells in the field of view (FOV) at the two-hour time point for each replicate. Cells were then categorised as either high, intermediate, or low phage-uptake using univariate clustering analysis (Fig. 1c and Supplemental Fig. 1) (46). A549 lung epithelial cells showed the highest accumulation of phages, with a median of cells containing fluorescently-labelled phages at the two-hour time point of 99% (± 2%, mean ± standard deviation [SD]; Field of View [FOV] *n* = 8; coefficient of variation [CV] = 2%). Next, HUVEC, MDCK-1, and HeLa cells, representing endothelial, fibroblast and epithelial cell types all showed intermediate levels of phage accumulation at the two hour time point, with medians of 44% (± 34% SD; FOV *n* = 10; CV = 63%), 51% (± 26% SD; FOV *n* = 4; CV = 54%), and 41% (± 18% SD; FOV *n* = 7; CV = 42%) of phage-positive cells, respectively. Finally, THP-1, HT29, and BJ cells, representing macrophages, epithelial and fibroblast all showed little to no accumulation with medians of 0% (± 11% SD; FOV *n* = 9; CV = 155%), 0% (± 0.5% SD; FOV *n* = 4; CV = 200%), and 0% (± 0% SD; FOV *n* = 4; CV = 0%) phage-positive cells at two hours, respectively. We further quantified the rate at which cells internalised phages by manually counting the number of cells per frame of interest containing fluorescently-labelled phages for each of the field of view per cell lines (Fig. 1d). Most cells showed large variability in the uptake rate over the two-hour period. For A549 cells, which had the highest accumulation of phages, we saw large variation in the rate of uptake, with a median of 27% (± 36% SD; FOV *n* = 6; CV = 83%) of cells that contained phages at one hour of incubation compared to 100% (± 0% SD; FOV *n* = 6; CV = 0%) of cells at two hours. Comparatively, HUVECs, which were classified as intermediate accumulation of phages showed extensive variability in their uptake rates, with a median of 12% (± 41% SD, FOV *n* = 8; CV = 116%) and 46% (± 35% SD; FOV *n* = 8; CV = 60%) of cells containing phages at one and two hours, respectively.

To confirm that phages were internalised and not simply attached to the cell surface, we created a three-dimensional (3D) reconstruction using a Z stack to visualise the intracellular localisation of phages. After the live cell imaging of MDCK-I cells incubated with fluorescently-labelled T4 phages for one hour (Supplemental movie SM8), we acquired a high-resolution Z stack image of one chosen field of view. We reconstituted the 3D volume of the cell to visualise phage repartition in the cytoplasm (Fig. 2 top left corner and Supplemental movie SM9). Finally, we looked at the localization of phages in 3D using the XY cross section (Fig. 2). The 3D reconstruction of the cell confirmed that phages internalized by the cells (visualised as green fluorescent particles) lie in the same focal plane as the nucleus (Fig. 2). Phages were distributed throughout the cell cytoplasm and appeared to be localised within membrane-bound vesicles surrounding the nucleus.

**Figure 2.**
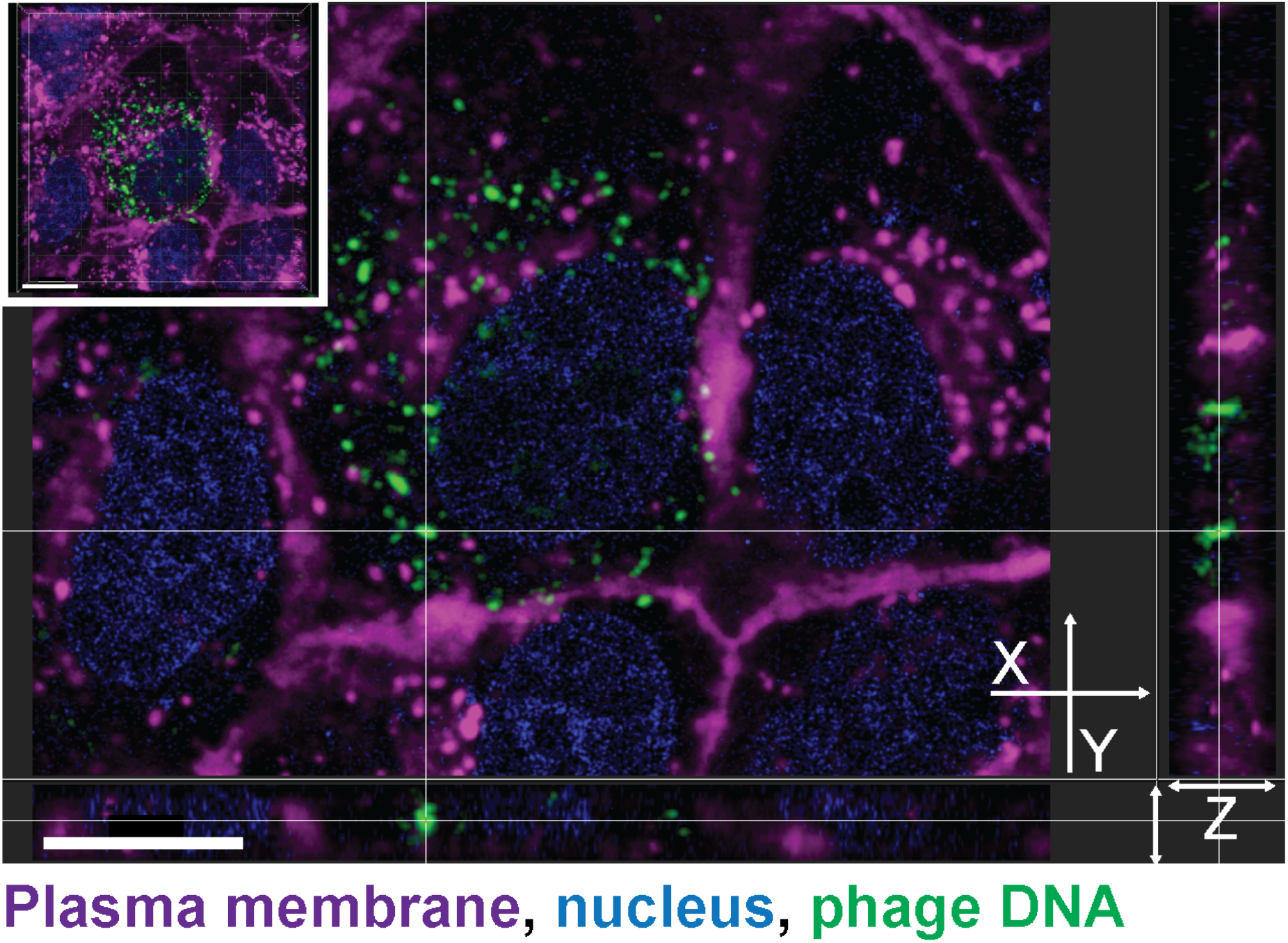
3D reconstruction of intracellular phages. MDCK-I cells were incubated for an hour with T4 phages on glass bottom slide before acquisition of a high-resolution Z stack to visualise phage dispersion inside of cells. Cells were stained with Hoechst 33342 nucleus stain (blue), CellMask plasma membrane stain (magenta) and T4 phages labelled with SYBR-Gold DNA-complexing stain (green). Images were acquired in real-time on live cells. XY cross section made using Imaris software with an embedded 3D projection of the cell in the top left corner of the image. The cross in the centre of the image shows a cluster of internalised phages with its repartition in the Z dimension (depth) of the image represented in the X and Y sides views. Scale bar: 10 µm.

### Phage uptake occurs at comparable rates under both static and flow conditions

The previous experiments were all conducted under static conditions, where phages were directly applied to the cell culture media and phage-cell encounters driven purely by diffusion. However, in the context of phage therapy, phages administered to the body are likely to encounter dynamic environments and active fluid flow, such as in the circulatory and lymphatic system. These dynamic conditions may lead to increased phage-cell encounter rates or altered cellular uptake (47–52). We investigated whether phage uptake rates under static conditions were comparable with uptake rates under fluid flow and shear conditions that mimic the circulatory systems of the body (Fig. 3a). We chose HUVECs to use in our flow experiment for two main reasons, the first one is that they are part of the intermediate uptake group determined in our first figure and are not part of one of the two other extremes. Secondly, HUVECs are endothelial cells and would be amongst the first type of cells to be in contact with circulating phages in the human body. Using an in-house fabricated microfluidic device mounted on a glass coverslip adapted for confocal microscopy (53,54), we seeded the device with HUVECs and incubated it under static conditions for 12 hours to ensure sufficient cellular attachment to the subtract. Cell layers were allowed to establish within the device under a low flow rate of 0.66 µl/min for one day, before increasing to a final flow rate of 8 µl/min until cells reached confluency. Physiological shear stress values observed in the human body ranges from 0.1 dyne/cm^2^ in the microcirculation, reaching higher rates of 50 dyne/cm^2^ found in larger circulatory vessels (55–59). Due to the volumes of media and quantity of phages applied to the chip, we chose a flow rate of 8 µl/min, which is equivalent to a shear stress of 0.1 dyne/cm^2^ in our chip and was at the lower end of physiological circulatory range (Supplemental Table 1). We perfused the chips with media containing 10^9^ phages/ml, with phage uptake visualised as previously described at two and 18 hours timepoints. Even though the volumes and quantity of phages seen by the cell layer in the static (200 µl) and flow (960 µl) conditions are different, we still observe similar rates of T4 phage uptake after two hours (Fig. 1a & 3b. ii and Supplemental Fig. 3). After 18 hours incubation under shear stress, we observed a significant increase in the fluorescence intensity compared with two hours incubation (unpaired t-test, P < 0.001) (Supplemental Fig. 3).

**Figure 3.**
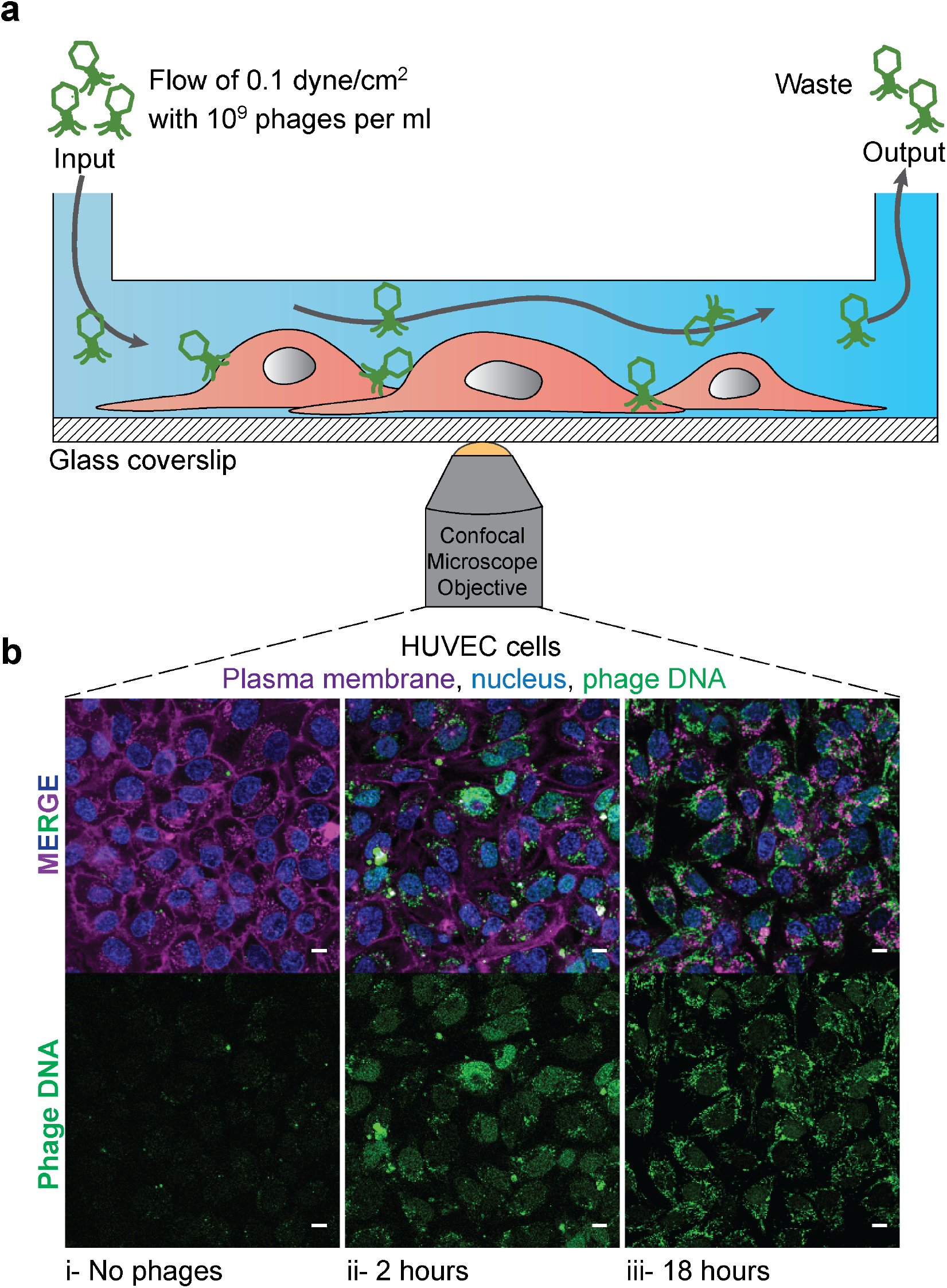
Uptake of phages under fluid flow and shear stress. T4 phage was applied to HUVECs within a microfluidic channel of a microfluidic device under a shear stress of 0.1 dyne/cm^2^ for two or 18 hours with images collected via real-time microscopy images. **(a)** Schematic of the microchannel showing the flow from one port of the channel to the other side of the channel where the waste was collected. **(b)** Cells were stained with nucleus stain (blue), plasma membrane stain (magenta) and T4 phages labelled with DNA-complexing stain (green). Control cells without phages i). Cells were incubated under a constant flow of phages at a rate of 8 µL/min equivalent to a shear stress of 0.1 dyne/cm^2^ for either ii) two or iii) 18 hours. Scale bar:10 µm.

### Phage size affects intracellular uptake

Next, we looked at the effect of phage size on the rate of cellular uptake. We chose three *Escherichia coli* infecting phages; T4 phage from the *Myoviridae* family, measuring 90 nm wide and 200 nm long with a contractile tail; Lambda phage from the *Siphoviridae* family, measuring 60 nm wide and 150 nm long with a non-contractile tail; and T3 phage from the *Podoviridae* family, measuring 55 nm wide and 30 nm long with a small non-contractile tail. We tested these phages against three cell lines representative of high, intermediate, and low rates of uptake (Fig. 1c); A549 cells with a high rate of uptake, HUVECs with an intermediate rate, and BJ cells with a low rate. We incubated phages with the cell layers for two hours, acquiring images every two minutes (Supplemental movies SM1-2, 7, 10-15), with the final time point represented in Fig. 4a. The first row of images shows clear differences in T4 phage uptake between the three cell lines (Fig. 4a). However, when we applied the smaller sized Lambda and T3 phages to the three cells lines, we saw a large increase in the uptake of both phages compared with T4. This was particularly evident in the BJ cell line, which had effectively no T4 phage update over a two-hour period but nonetheless, demonstrated increased uptake of the smaller sized Lambda and T3 phage. We quantified phage uptake across phage size and cell type using a pipeline built with CellProfiler software (60) (see methods) to analyse the median grayvalue intensity in the cell region (median over all pixels in FOV marked as cells) as a proxy for fluorescence intensity of phage (Figure 4). For T3 phage, the smallest phage of the three tested, we observed the highest rate of uptake across the three cell lines (ANOVA one way, F (2, 45) = 71,32; P < 0.0001). This was especially evident for the BJ cell line where the median intensity of the phage fluorescence signal increased from a median of 0.01 normalised grey value with T4 phage (± 0.006 SD; FOV *n* = 6; CV = 49%) up to 0.09 with Lambda (± 0.04 SD; FOV *n* = 6; CV = 49%), and finally to 0.16 with T3 phage (± 0.04 SD; FOV *n* = 3; CV = 22%) (Fig. 4b). Based on our microscopy results, we suggest that smaller sized phages broadly increase the rate of cellular uptake, and that these effects were more pronounced in our intermediate and low uptake cell lines.

**Figure 4.**
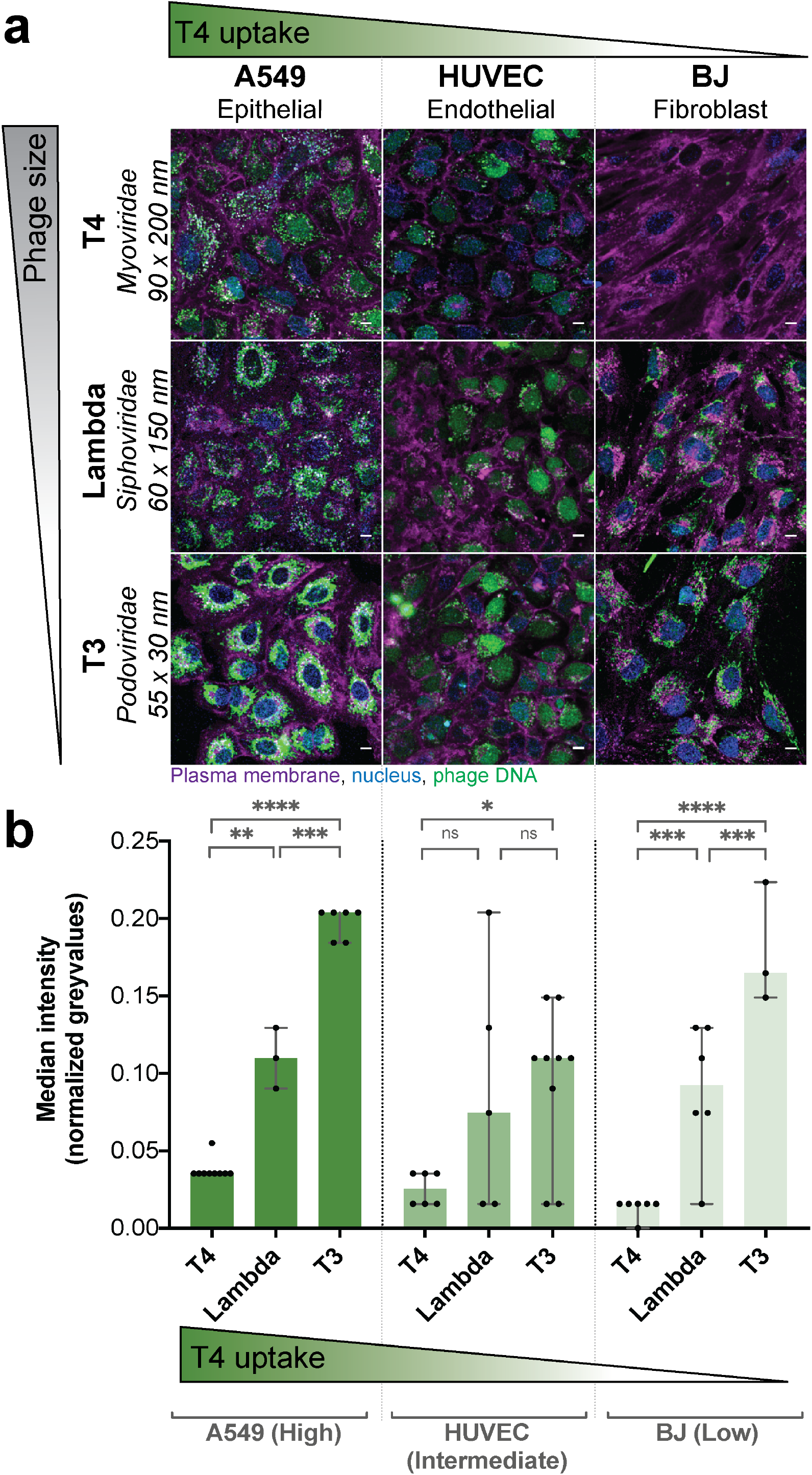
Cellular uptake of differing phage particle sizes. Real-time microscopy images showing differential uptake of phages based on particle size. One representative cell line from each of the three high, intermediate, and low uptake groups were picked. **(a)** Images were acquired in real-time. Cells were stained with Hoechst 33342 nucleus stain (blue), CellMask plasma membrane stain (magenta) and T4 phages labelled with SYBR-Gold DNA-complexing stain (green). Three cell lines used; A549, HUVEC and BJ, with the three phages; T4, Lambda, and T3. Green gradient shows the qualitative increase in T4 phage uptake shown in Figure 1C. Grey gradient shows qualitative decrease in sizes of phages T4 (90 × 200 nm), Lambda (60 × 150 nm), and T3 (55 × 30 nm). Scale bar = 10 µm. The image shown is the last image at two-hour acquisition. **(b)** Fluorescence intensity of the “phage object” area in normalized grey values quantified from the CellProfiler software of the phage channel fluorescence at the two-hour time point (A549 – T4 *n* = 9; A549 – Lambda *n* = 3; A549 – T3 *n* = 6; HUVEC – T4 *n* = 6; HUVEC – Lambda *n* = 3; HUVEC – T3 *n* = 6; BJ – T4 *n* = 6; BJ – Lambda *n* = 6; BJ – T3 *n* = 3). Scatter plots show medians over all pixels in FOV marked as cells; error bars represent 95% confidence intervals; each point represents one FOV. P-values calculated from a one-way ANOVA (P < 0.0001: ****; P < 0.05: *; ns: non-significant).

### Phages rapidly adhere to eukaryotic cells, resulting in inactivation and uptake

We demonstrate that phages have different rates of cellular uptake depending on both cell type and the size of phages. Yet whether these phages remained functional, or if they were inactivated by the cellular uptake and trafficking pathways remains unclear. To answer this, we quantified the number intracellular phages using two methods; classic plaque formation unit (PFU) assays and Droplet Digital PCR (ddPCR). Briefly, PFUs allowed us to quantify the number of active or functional phages present within the cells, while ddPCR quantified the absolute number of phage DNA genome copies present in the sample. We evaluated the accuracy of the two techniques with a dilution series of our initial phage sample from 10^9^ to 10^2^ phages per ml using PFU and ddPCR (Supplemental Fig. 4). Subtracting active phages (PFU) from phage DNA genome copies (ddPCR) enabled us to quantify the proportion of phages inactivated or damaged during cellular uptake and trafficking, as these damaged phages are no longer able to infect their bacterial host and therefore will only be detected by ddPCR.

In order to differentiate between intracellular phages and those adhered to the cell surface, we incubated cells with T4 phages (∼10^9^ per ml) for two different periods of time: 30 seconds, which is too short for phage internalisation; and 18 hours, to maximise the number of internalised phages. After incubation at each time point, cells were extensively washed to remove non-adherent phages, lysed, and phages quantified by both PFU and ddPCR (Fig. 5a). We compared the same representative cell types previously used for high, intermediate, and low uptake cell lines (Fig. 5b). For all three cell lines, we saw between 90 and 2200 active phages (PFU) per ml adhered to the cells within 30 second treatment, with a median of 625 phages/ml (± 420 SD; *n* = 6; CV = 60.1%), 325 phages/ml (± 300 SD; *n* = 6; CV = 76.6%) and 250 phages/ml (± 850 SD; *n* = 6; CV = 114%) for A549, HUVEC and BJ cells, respectively, suggesting a small, yet persistent number of phages rapidly adhere to the cell layers. After 18 hours of incubation, we saw a large increase in the number of functional phages associated with the cells, with between 1.1 × 10^4^ to 3.1 × 10^6^ phages per ml accumulating within the three cell lines. Looking across the different cell lines, we see the highest accumulation of phages in the intermediate uptake cells, with a median of 5 × 10^5^ phages/ml (± 6.4 × 10^5^ SD; *n* = 9; CV = 85.7%) followed by the high and low uptake with medians respectively of 1.2 × 10^5^ phages/ml (± 8.4 × 10^4^ SD; *n* = 9; CV = 65.7%) and 5 × 10^4^ phages/ml (± 1.4 × 10^6^ SD; *n* = 9; CV = 144.3%), although there were no significant difference between the three cell lines (P > 0.1, ANOVA one way). We note that the BJ cell line, which showed the lowest rate of uptake observed via microscopy (Figure 1), still accumulated active phages over prolonged periods of time with non-significant differences of active phages at 18 hours observed with two other cell lines (unpaired t-test, two tailed, P > 0.08 with A549 and P > 0.6 with HUVEC).

**Figure 5.**
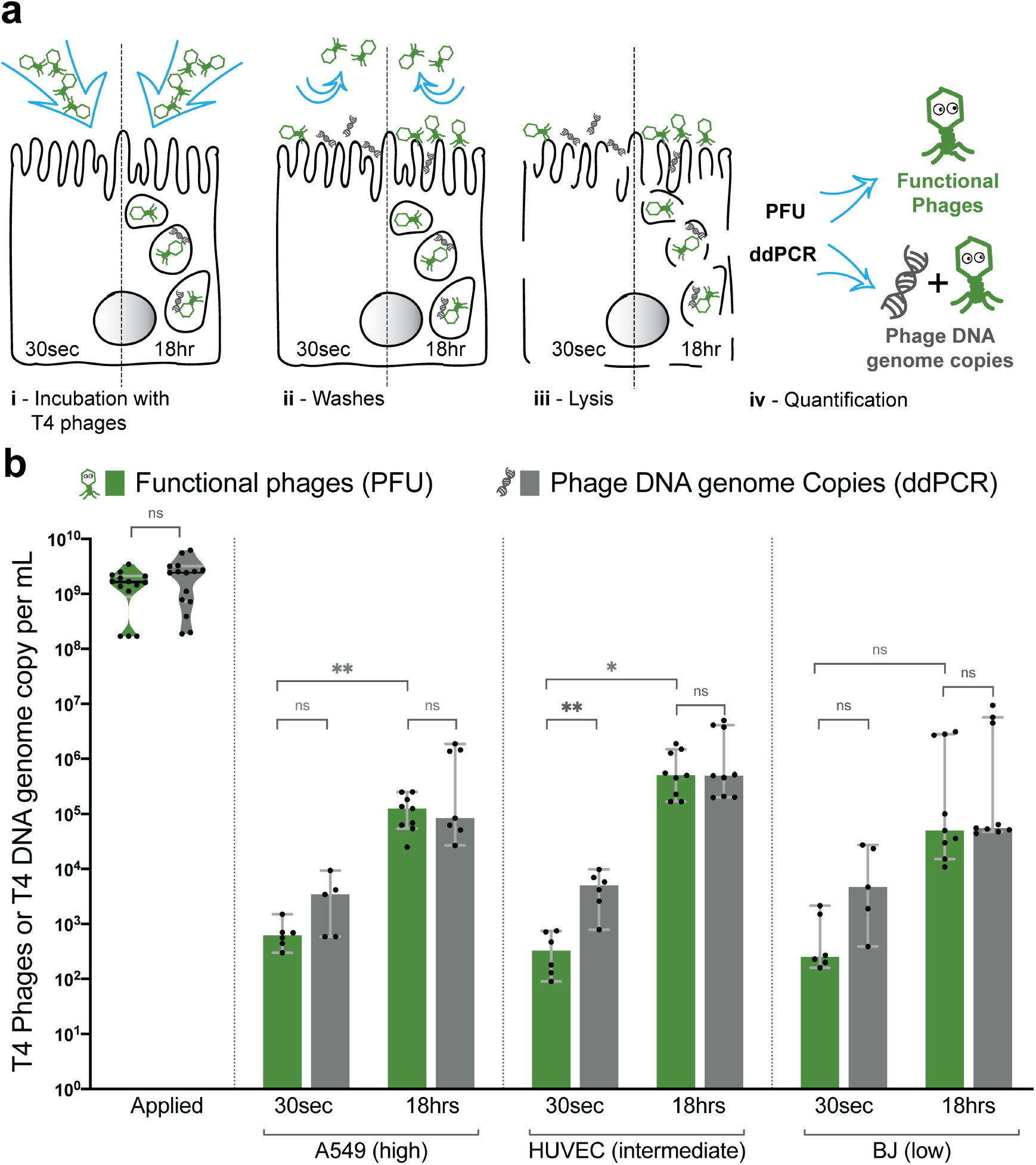
Quantifying adherence and internalisation of phages and their activity. Applied T4 phages were quantified using both traditional plaque assays (PFUs) and Digital Droplet PCR (ddPCR), across three different cell-lines: A549, HUVEC and BJ, representative of the high, intermediate, or low phage uptake. Phages were applied to cells for either 30 seconds as an adherence control, or for 18 hours to saturate phage uptake. (**a**) Schematic representation of the experiment showing steps taken to obtain the samples with two incubation times: 30 seconds or 18 hours. Active or functional phages are represented in green and phage DNA or inactive phages are represented in grey; (i) Incubation of phages with cells, (ii) Extensive washing of cells to remove non-adherent phages, (iii) Chemical and mechanical cell lysis, and (iv) Quantification of the active and total phages via PFU and ddPCR. (**b**) Active T4 phages in green quantified by PFU and in grey the total number of T4 phage DNA genome copies per sample quantified by ddPCR, including both active and inactive phages. Scatter plots show medians of phages or DNA genome copies per mL; error bars represent 95% confidence intervals; each point represents one sample. P-values calculated from an unpaired t-test between each pair (P < 0.001: ***; P < 0.01: **; P < 0.05: *; ns: non-significant).

Surprisingly, when looking at ddPCR results, we see an increase in phage DNA associated with the 30 second treatment in all cell lines, with between 3.9 × 10^2^ and 2.7 × 10^4^ DNA genome copies per ml persisting. When quantifying the inactivated phages at 30 second treatment, which was calculated as the difference in DNA genome copies and PFU, we observe between 7 × 10^1^ and 2.7 × 10^4^ inactivated phages per ml. These results suggest that; 1) phages rapidly adhere to eukaryotic cell layers, with a portion of these phages being inactivated, 2) longer incubation time allows for adhered phages to be internalised and accumulate inside of the cells, and 3) that the majority of internalised phage particles remain active and stably persist within the cells for up to 18 hours.

### Phage inactivation and internalisation influences pharmacokinetics

A key underexplored aspect of phage pharmacokinetics (PK) are the non-specific interactions between phages and the epithelial and endothelial cell layers of the body. To address this, we integrated our experimental data (Fig. 5) with an established PK model for phage administration in a rats (38). The model was established from a single-dose PK study performed in healthy rats, following an intravenous bolus of phages. The phage disposition in blood was well characterised using a standard three-compartment PK model (38). In order to evaluate the impact of phage inactivation and internalisation by eukaryotic cells on phage distribution, a fourth-compartment was added to the existing model, representing the epi- and endothelial cell layers (Fig. 6a). Deterministic simulations were subsequently performed with the previously estimated PK parameters (38) and the first-order inactivation constant estimated from our *in vitro* data (ddPCR). Inactivation rate constant was calculated using both the 30 seconds- and 18 hours-derived constants, as a representative of both the rapid and prolonged phage accumulation by cell layers (Fig. 6b, supplemental data SD2, SD3 and table 5). Deterministic simulations were performed at an IV bolus dose of 10^9^ phages. With the inactivation rate constant calculated using 30 second data, complete removal of phages was noted as short as three hours. This is consistent with the 30 second *in vitro* data, in which rapid inactivation was observed. Comparatively, using the inactivation rate constant calculated using the 18 hours data, an initial rapid decay of circulating phages, followed by a longer tail and phage persistence in the blood was observed for up to 32 hours. These two models represent two extremes of rapid and prolonged phage accumulation by cell layers, highlighting the potential impact of these cellular mechanisms on phage disposition. Further studies characterising both the affinity of phage– eukaryotic interactions and their influence on PK are needed for better clinical translation.

**Figure 6.**
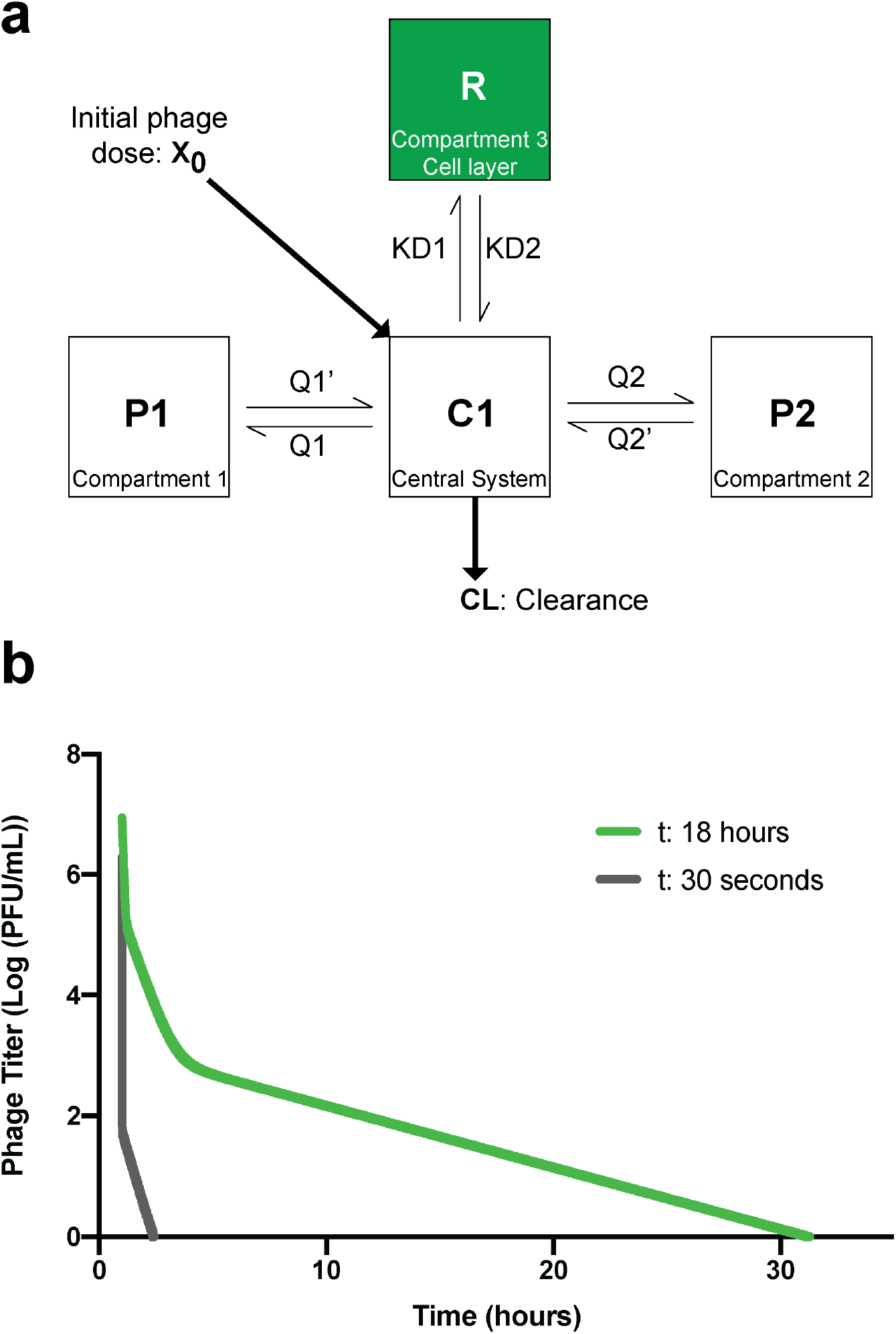
Pharmacokinetic (PK) model of inactivation and internalisation of phages by eukaryotic cell layers. Extrapolation of *in vitro* results were fit to a rat model of phage decay at an initial dose of 10^9^ phages. **(a)** Schematic of the standard 3-compartments model with an additional inactivation compartment. X_0_, the initial dose of phages and CL phage total body clearance. C1, the central compartment representing the central system (*i.e*., the blood). P1 and P2 compartments represent various organs or tissues participating in phage decay. R, the new compartment representing *in vitro* cell layers as a major sink of phages. Q1 and Q2 are intercompartmental clearance one and two, respectively. KD1 is the first order inactivation constant and KD2 is the first order reactivation constant (assumed to be zero in our model). **(b)** Graph representing the decay of phages per central volume of distribution over time calculated with *in vitro* data using an R model with an integration step of 0.001. First order for the 30 seconds graph is of 1511 1/h/rat and for the 18 hours the first order is of 0.44 1/h/rat calculated from the geometric mean of ddPCR data (calculation in supplemental table 5).

## Discussion

Phage therapy is one of the more promising solutions in the fight against antibiotic resistance (25). However, the effective use of phages as a treatment against multi-drug resistant bacterial pathogens remains a major challenge to successfully and reproducibly implement. Our understanding of the interactions between bacteriophages and eukaryotic cells may have important implications and impacts for the success of phage therapy. In this article, we investigate the interactions between phages and *in vitro* tissue culture cell layers. We demonstrate that cell type plays a major role in the uptake process of phages, with individual cells taking up phages at different rates, even amongst the same cell types (e.g., epithelial, endothelial, etc.). Uptake rates were validated under shear stress conditions using a microfluidic device that mimics the shear stress to which endothelial cells are exposed to *in vivo* (61,62), and shown to be comparable with static conditions. Using live cell imaging we show that phages accumulate within cells over time and that smaller phages are internalised at higher rates than larger phages. Phages were found to rapidly adhere to eukaryotic cells, with adherent phages being internalised by macropinocytosis over the following 18 hours of incubation, with functional phages accumulating and stably persisting within the cells. Finally, we incorporate our results into an established pharmacokinetic (PK) model (38), demonstrating the potential impact of phage accumulation by epithelial and endothelial cell layers, which represent an unaccounted sink of phages within the body.

The mechanism of non-specific phage uptake by *in vitro* tissue culture cell lines was previously demonstrated to occur via macropinocytosis (8). Cell types have been reported to have differing basal rates of macropinocytosis depending on their function and location within the body (24), thereby influencing their rate of phage uptake. The macropinosome plays further roles in the presentation of antigens for pattern recognition receptors located in other organelles, and in the activation of innate immune responses (63). The fate of phages within the macropinosome is still unknown. It is clear phages accumulate within the macropinosomes over time, but whether they are subsequently degraded by the endosomal/lysosomal system, or recycled and fused back with the plasma membrane remains to be investigated (24). Here we propose that, within seconds, phages rapidly adhere to the eukaryotic cell membrane. Secondly, this adherence leads to an internalisation of phages via non-specific macropinocytosis. Thirdly, this allows for the steady accumulation of phages within the macropinosomes and cell over time. Further research is required to probe the specific mechanisms of phage adherence, uptake, and the cellular mechanisms that govern the trafficking of phages within cells (64,65).

Using three different phages, from different families, with different sizes, we were able to show an effect of phage size on the rate of uptake across three different cell lines. Using real-time microscopy, we demonstrate that the smaller sized phage particles had increased rates of uptake, especially in low uptake cell lines. One hypothesis for the increased uptake of smaller sized phages, may simply be the result of an increased number of particles capable of interacting with actin-mediated ruffles associated with of macropinosome formation, thereby leading to a higher number of phages engulfed within each macropinosome (24,66). It is also possible that there is a difference in ruffle size across cell lines leading to a predisposition of some cells to uptake smaller or bigger size phages. A similar relationship between particle size and uptake has been made with nanoparticles, where it was observed that 50 nm size nanoparticles had high rates of uptake, which coincidentally are of similar size as our T3 phages (67–69). It was also shown that the cell type, as well as the particles shape influenced uptake, with disc-shaped particles having higher rates of uptake compared with elongated rod shapes. Again, our T3 phage are coincidentally similarly shaped to the disc-shaped particles (70,71), indicating that the shape of phages may play a role in their uptake; a factor that we cannot rule out from our study. It is intriguing to speculate that much of the research into nanoparticle delivery has converged upon parameters analogous with biology’s own naturally-occurring nanoparticles; the bacteriophages.

We observed, that following the rapid adsorption of phages with *in vitro* cell culture layers, a high proportion of phages were inactivated (Fig. 5b); a phenomenon which has not been previously reported. Our first hypothesis to explain this rapid inactivation is that upon interaction with cell surface features, phages are mechanically triggered to eject their genomes, thereby leading to inactivation (72). This would implicate an increase of phage DNA at the cell surface, which we observed via ddPCR results, along with a concomitant decrease in functional phage particles. Another hypothesis is that the transient and non-specific interactions between the phages and the cell surface features, including glycoproteins, glycolipids, and mucins (73), may physically block or impede the phage such that it is unable to access its host bacterial receptor for plaque quantification, thereby only being detected via ddPCR. Finally, this inactivation could be the result of the degradation of the phage capsid due to enzymes, secretions, or cellular products. Though our negative controls show this is unlikely, as there was no effect of spent cell culture media on phage infectivity (Supplemental Fig. 5). At this stage whilst we cannot conclude the specific mechanisms inducing this rapid loss of phage viability, our results clearly suggest that this rapid inactivation and adsorption to the cell layer may represent a major sink for circulating phages in the body.

During phage therapy, the cells, organs, and systems of the body play an important role in the efficacy of treatment due to their effect on the sequestration of active phages and limiting accessibility to the site of infection (30,31). It has been proposed that the mononuclear phagocyte system (MPS) is primarily responsible for the filtering and removal of the phages during phage therapy, with the liver and spleen considered the main organs responsible for filtering circulating phages (43,74,75). Recent case studies of phage pharmacokinetics (PK) following intravenous (IV) administration have shown rapid clearance in both humans and animal models, with over > 99% of phages applied removed from circulation within the first few hours (38,76). In an extensive literature review, Dabrowska 2019 (30) noted that the phage titres in the blood immediately following intravenous injection (1-5 minutes) were markedly less than expected hypothetical dilutions. Even when accounting for phage dilutions in the blood or total body volume, phage titres only reached between 0.02% and 0.4% of their predicted titres. This suggests that there is significant and rapid uptake of phages by the organs and cells of the MPS, or alternatively, the rapid adherence and inactivation of phages to the epi- and endothelial cells lining the circulatory system. All the cell lines used in this study; endothelial, epithelial, macrophages or fibroblast, may be in contact with phages at any time during therapy. Our results demonstrate that phages rapidly adhered to and are subsequently internalised by these cells (Fig. 5b). The model we developed in this article (Fig. 6), whilst preliminary, illustrated the potential impact that cell layers play in the inactivation of phages following their delivery to a patient. We suggest that the cell layers of the body are a major sink for phages and that these interactions have unrecognised impacts on phage PK and therapy. If this mechanism is true, then phage most likely display non-linear PK (in addition to non-linear clearance). These results have unrecognised consequences on phage dosing regimens during treatment, implying higher doses and more frequent administrations of phages to patients may be needed in order to reach an optimal phage dose to fight the infection.

With these new findings on the role of eukaryotic cells in the uptake and inactivation of phages during phage therapy, we hope to help in the design and engineering of treatment for patients and to improve the clinical outcome of phage therapy.

## Materials and methods

### Bacterial stocks and phage stocks

The bacterial strains used in this study, *Escherichia coli B* strain HER 1024 and *E. coli B* strain W3350, were cultured in lysogeny broth (LB) media (10 g tryptone, 5 g yeast extract, 10 g NaCl, in 1 litre of distilled water [dH_2_O]) at 37 °C shaking overnight and used to propagate and titre phages T4, T3 and lambda supplemented with 10 mM CaCl_2_ and MgSO_4_. Phages T4, T3 and lambda were cleaned and purified using the Phage on Tap protocol (PoT) (45) and titred up to a concentration of approximately 10^10^ phages per ml. After purification, phages were stored in a final solution of SM Buffer (2.0 g MgSO_4_·7H_2_O, 5.8 g NaCl, 50 ml of 1M Tris-HCl pH 7.4, dissolve in 1 litre of dH_2_O) at 4 °C.

### Endotoxin removal

For each of the phage samples, endotoxin removal protocol was followed from the Phage on Tap (PoT) protocol (45). Phages lysates were cleaned four times with octanol to remove endotoxins from the lysate.

### Cell line stocks

Seven cell lines were used in this study, all grown at 37 °C and 5% CO_2_ and supplemented with 1% penicillin-streptomycin (Life Technologies Australia Pty. Ltd) A549 cells were grown in Ham’s F-12K (Kaighn’s) (also called F12-K)) (Life Technologies Australia Pty. Ltd) media with 10% Fetal Bovine Serum (FBS) (Life Technologies Australia Pty. Ltd), HUVECs were grown in F12-K media with 20% FBS, HeLa and HT-29 cells were both grown in Dulbecco’s Modified Eagle Medium (DMEM) (Life Technologies Australia Pty. Ltd) supplemented with 10% FBS, MDCK-I cells were grown in Modified Eagle Medium (MEM) (Life Technologies Australia Pty. Ltd) with 10% FBS, BJ cells were grown in DMEM media with 10% FBS and 1% sodium pyruvate (Sigma-Aldrich, Australia) and finally the suspension of THP-1 cells were maintained in Roswell Park Memorial Institute (RPMI) 1640 media (Life Technologies Australia Pty. Ltd) with 10% FBS. For differentiation, phorbol 12-myristate 13-acetate (PMA) (Sigma-Aldrich, Australia) was added to a final concentration of 25 mM and incubated for 48 hours. After incubation PMA supplemented media was removed and cells were further grown in PMA free media for 24 hours to obtain differentiated macrophages. These differentiated cells were stable for up to one week.

### Confocal microscopy

For confocal microscopy experiment, cells were seeded in an IBIDI μ-Slide 8-well glass-bottom slide (DKSH Australia Pty. Ltd) and grown to 80-90% confluency for acquisition. Cells were treated for 20 min with the respective culture media for each cell line with 5% Hoechst 33342 stain, excitation/emission 361/497 nm (Life Technologies Australia Pty. Ltd) and 1% CellMask deep red plasma membrane stain, excitation/emission 649/666 nm (Life Technologies Australia Pty. Ltd). After incubation cells were washed three times with Dulbecco’s phosphate-buffered saline (DPBS) 1× and then left in Hank’s Balanced Salt Solution (HBSS) with 1% FBS until acquisition. Purified phages were labelled with 1% SYBR-Gold, excitation/emission 495/537 nm (Life Technologies Australia Pty. Ltd) for one hour in the dark at 4 °C, followed by three washes with HBSS in Amicon-Ultra4 centrifugal unit 100-kDa membrane (Merck Pty. Ltd) to remove excess of stain. The washed phages were resuspended in a final volume of 1 ml in HBSS media. From a 10^9^ phage per ml solution, we added 200 μl per well to the cells under the microscope right before the start of the acquisition. Cells were imaged with HC PL APO 63x/1.40 Oil CS2 oil immersion objective by Leica SP8 confocal microscope on inverted stand with a hybrid detector (HyD) in real time. Excitation used for Hoechst 33342 (blue), SYBR-Gold and CellMask deep red was 405, 488 and 638 nm; corresponding emission was recorder at 412-462, 508-545 and 648-694 nm detection ranges respectively. HyD detector was used in sequential mode to detect the phages, it increases the sensitivity of detection by acquiring the same image multiple times and accumulating the fluorescence signal. All live cell imaging experiments were completed in triplicate (three fields of view per session). One image was acquired every two minutes for two hours. Each field of view was hand-picked depending on the cell confluency and success of staining. Videos were created through post-processing using the FIJI software version 2.0.0-rc-68/1.52f (77). First, the three channels acquired were merged and processed with a Gaussian Blur filter of 0.8. Second, each channel brightness and contrast were enhanced for printing quality. Finally, the time and scale were added to the final movie saved in 12 fps.

### Quantification of phages in live cell imaging

For each live cell experiment, we quantified cells that contained intracellular green fluorescence as indicative of SYBR-Gold labelled phages. Live cell images were acquired every ten minutes were quantified by manual counting the total number of cells in the field of view and the number of cells with intracellular phages to calculate the percentage of cells containing intracellular phages. Results were plotted using the GraphPad Prism version 8.4.2 for macOS GraphPad Software, San Diego, California USA, www.graphpad.com, to show uptake of phages over time.

### Clustering analysis

Univariate clustering was performed using the dynamic programming algorithm in the R package Ckmeans.1d.dp (46).

### Flow conditions in microfluidic chip

A chip mold with 500 µm wide, 350 µm high and 1.3 cm long channels was designed using SolidWorks® 2017. The moulds were 3D-printed using Object Eden360V (Stratasys, USA) with a manufacturer-provided polymer FC720 and surface-salinized in a vacuum desiccator overnight with 20 µl Trichloro(1*H*,1*H*,2*H*,2*H*-perfluorooctyl)silane (Sigma-Aldrich, USA). The microfluidic chips were manufactured via soft-lithography, by casting a 10:1 mixture of Sylgard 184 PDMS and its curing agent (Dowsil, USA) respectively, onto the moulds and were cured at 90 °C until completely solidified (∼2 hours). The chips were then cut with a surgical knife, gently peeled off, trimmed and their inlet and outlet were punched with 2 mm biopsy punchers (ProSciTech, Australia). Subsequently, the chips were washed in pentane and acetone to remove residual uncured PDMS. Atmospheric plasma generated at 0.65 Torr with high radio frequency was used to bound the PDMS chip to a glass cover slip No. 1.5H (0.170 mm ± 0.005 mm thickness) optimized for confocal microscopy (Marienfeld), for 20 seconds. The microchannel of the assembled microfluidic device were washed with ethanol (80% v/v)-sterilised, UV-sterilised and pre-treated with 1:50 MaxGel™ ECM (Sigma-Aldrich) in cold F12-K media for two hours at 37 °C and 5% CO2. The microchannel was then washed with F12-K media to remove residual ECM. Schematic and picture of the microdevice is included in Supplemental Fig. 2. The channel was seeded with 10 µl of HUVECs at a concentration of 5 × 10^5^ cells/ml, which were carefully pipetted through the in port. The seeded chip was incubated statically for 12 hours to allow cell attachment at 37 °C and 5% CO2. This was followed by perfusing the attached cells with complete media for 24 hours at 0.66 µl/min flow rate to establish a confluent cell layer. The cell layer was then perfused with cell culture media supplemented with 20% of FBS for another 24 hr at 8 µl/min to acclimate the cells to the shear stress. Perfusion was mediated by a single-channel syringe pump (New Era Pump Systems, USA) using a 10 ml 21 gauge needled-syringe fitted to Teflon tubes of 1/16” inner diameter and 1/32” outer diameter (Cole-Palmer, USA) that were previously sterilised using ethanol (80% v/v)-sterilised, DPBS and UVs. HUVECs were then stained with nucleus stain, Hoechst 33342 (blue), plasma membrane stain, CellMask (magenta) under static conditions for 20 min. T4 phages labelled with DNA-complexing stain, SYBR-Gold (green) were then added to the chip under 8 µl/min flow rate for either two or 18 hours. After incubation under flow with the phages, the in and out port of the chips were sealed using sterilized-binder paper clips and the chip placed under the microscope. The images were acquired with HC PL APO 63x/1.40 Oil CS2 oil immersion objective on an inverted Leica SP8 confocal microscope. A hybrid detector (HyD) was used to visualise phage DNA (excitation/emission 495/537 nm), other channels were acquired with conventional PMT detectors for CellMask (excitation/emission 649/666 nm) and for Hoescht 33342 (excitation/emission 361/497 nm).

### Image analysis with CellProfiler

To quantify the fluorescence intensity of SYBR-Gold labelled phages (495 nm wavelength), we used a pipeline created in CellProfiler (60) (see the pipeline used in supplemental data, SD1), allowing us to measure the pixel grey values as a proxy for fluorescence intensity across the image. First, we segmented regions covered by nuclei by applying the IdentifyPrimaryObjects module to the Hoechst channel image. Second, we defined expanded regions around the nuclei for cytoplasmic measurements using the IdentifySecondaryModule with the parameter Distance-N set to 200. Third, we masked out nuclei regions in the of the nuclei SYBR (phages) channel. This is to exclude fluorescence coming from the cell nuclei due to the leaking of the SYBR dye from the phage capsid to the cell nuclei, which would lead to false positive quantification. Finally, the grey values image intensity in the masked SYBR channel and additional parameters of the secondary objects were measured (Supplemental data SD1). Only a single time point at two hours at each field of view was used for the analysis. The number of images analysed for each condition varied, as manual quality control was applied to exclude out of focus and non-analysable fields of view.

### Intracellular phages

For the intracellular phages experiment, cells were plated in T25 cm^2^ flasks until they reached confluency. For the 18 hours experiment, phages were applied in volumes of 3 ml of media with 10^9^ phages/ml per flask and incubated overnight at 37 °C and 5% CO_2_. The control flasks were incubated with 3 ml of phage-free media. After the 18 hours incubation, control flasks were incubated with the same phage dilution for 30 seconds. The initial dilution for each flask was collected for quantification. Cells were washed three times with 5 ml of 1 × DPBS to remove non-adherent phages. Next, one ml of 0.5% trypsin was added per flask and incubated at 37 °C and 5% CO_2_ for a few minutes, once cells detached, the cells were resuspended in 5 ml of 1 × DPBS and spun at 1500 rpm for three minutes and washed three times with 5 ml 1 × DPBS to remove any non-adherent phages. After the washes, cells were resuspended in 1 ml of lysis buffer (0.5 M EDTA and 1 M Tris at pH 7.5, complete with dH_2_O and adjust pH to 8) and left at room temperature for 20 min. After incubation the cells are passed through a 30 G syringe three times to ensure complete cell lysis. The lysis was confirmed by looking at the sample under a microscope.

### ddPCR setup

Digital Droplet Polymerase Chain Reaction (ddPCR) was performed following manufacturer’s instructions (Bio-Rad, Australia). A 20 µl reaction was assembled with primers, probe, ddPCR Supermix for probe (Bio-Rad, Australia) and sample. The primer and probe sequence and PCR parameters are shown in supplemental Table 2 – 4. ddPCR reaction mix was then loaded into eight channel disposable droplet generator cartridge (Bio-Rad, Australia). 70 µl of droplet generation oil was added to each channel and placed in the Bio-Rad QX200 droplet generator. The droplets were transferred into the deep well 96 well plate (Bio-Rad, Australia), using a multichannel pipette. The plate was then sealed using Bio-Rad plate sealer and then placed in a conventional thermocycler and the PCR product was amplified (supplemental Table 4). After amplification, the plate was loaded into the droplet reader (Bio-Rad, Australia) to quantify the fluorescent droplets. Analysis of the data was performed using the Poisson distribution with QuantaLife software (Bio-Rad, Australia).

**PFU quantification**. The Plaque Forming Unit (PFU) assay was performed using LB agar plates where a thin layer of soft LB agar was mixed with one ml of host bacterial culture and the desired dilution of phages was poured on the agar plate. The plate was incubated overnight at 37 °C before counting the number of plaques formed on the bacterial lawn. The results were calculated in PFU. To obtain the number of inactive phages we subtracted PFU numbers (active phages) from the ddPCR numbers (total number of phages).

### Pharmacokinetics Model

A previously developed PK model in healthy rats was utilized to evaluate the impact of phage inactivation on *in vivo* phage disposition (38). An additional compartment was incorporated to describe the inactivation and reactivation of phages by the epi- and endothelial cells. The rates of inactivation and reactivation was described by first-order rate constant, KD, and was assumed to be constant over time. The differential equations for phage disposition and inactivation were represented by:

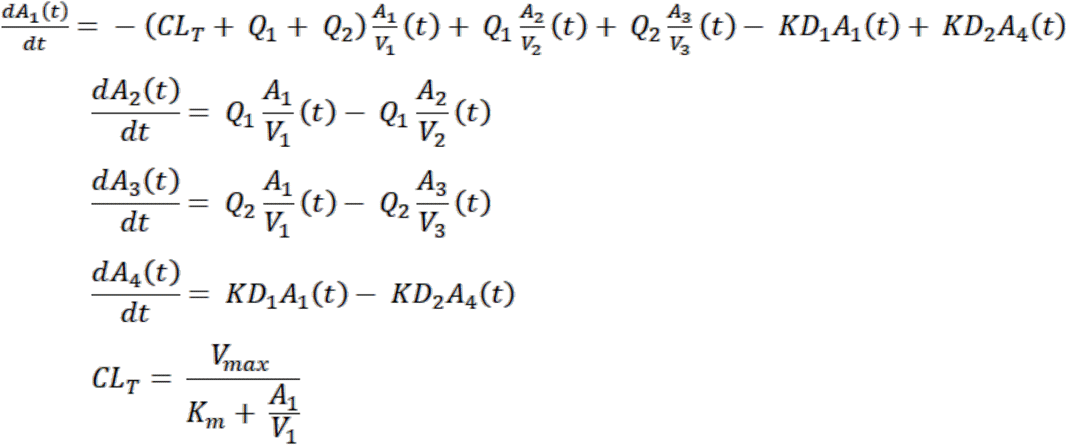

where

*Q*_1_= inter-compartmental clearance 1 (mL/h/Rat).

*Q*_2_= inter-compartmental clearance 2 (mL/h/Rat).

*V*_1_= Volume of distribution of the central compartment (mL/Rat).

*V*_2_= Volume of distribution of the peripheral compartment 1 (mL/Rat).

*V*_3_= Volume of distribution of the peripheral compartment 2 (mL/Rat).

*k*_*m*_= Phage titre that produces 50% of the maximal elimination rate of the system (PFU/mL/Rat).

*V*_*max*_= Maximum elimination rate (PFU/h/Rat).

*KD*_1_= Inactivation rate constant (1/h).

*KD*_2_= Reactivation rate constant (1/h).

Deterministic was performed using model-predicted median PK parameters in rats without inter-individual variability and random unexplained variability (Supplementary table 5). Inactivation rate constant was determined using the ddPCR results as described in Supplementary table 5. First order for the 30 seconds graph is of 1415 1/h/rat and for the 18 hours the first order is of 0.358 1/h/rat calculated from the ddPCR data. Reactivation rate constant was fixed to 0. Deterministic simulations were performed in R using mrgsolve (version 0.10.4) (38,78).

## Supporting information

Supplemental Movie 02_HUVEC-T4

Supplemental Movie 03_MDCK-I-T4

Supplemental Movie 04_HeLa-T4

Supplemental Movie 05_THP-1-T4

Supplemental Movie 08_MDCK-I-T4

Supplemental Movie 09_MDCK-I-T4-3D

Supplemental Movie 10_A549-Lambda

Supplemental Movie 11_HUVEC-Lambda

Supplemental Movie 12_BJ-Lambda

Supplemental Movie 14_HUVEC-T3

Supplemental Data 1_CellProfilerProject

Supplemental Data 2_RServer

Supplemental Data 3_Rui

## Authors participation

Conceptualization: MCB, JJB; Methodology: MCB, WHC, WR, AO, LAF, CAH, YWL; Formal Analysis: MCB, JJB; Investigation: MCB; Resources: JJB; Writing – Original Draft Preparation: MCB, RP, JJB; Writing – Review and Editing: all authors contributed; Supervision and Funding Acquisition: JJB

## Acknowledgements

We thank the following labs for kindly providing the cell lines; Hudson Institute of Medical Research and the Oncogenic Signalling Lab for providing the A549 cell line; the Nucleic Acids and Innate Immunity Research Group for providing the HT29 and BJ cell lines; the University of Melbourne and the Obstetrics, Nutrition and Endocrinology Group for providing the HUVECs; Monash University and the Moseley Laboratory for providing the HeLa cell line. We thank the following facilities for kindly providing equipment and guidance; Monash Micro Imaging facility for help with microscopy acquisition, Monash School of Engineering for providing support in the fabrication of microfluidic devices, and Department of Biochemistry & Molecular Biology (Monash Biomedicine Discovery Institute) Imaging Facility for providing access to the ddPCR equipment. This work was performed in part at the Melbourne Centre for Nanofabrication (MCN) in the Victorian Node of the Australian National Fabrication Facility (ANFF).

## References

1. Rohwer F, Segall AM. In retrospect: A century of phage lessons. Nature. 2015;528(7580):46–8.

2. Hatfull GF. Dark Matter of the Biosphere: the Amazing World of Bacteriophage Diversity. J Virol. 2015 Aug;89(16):8107–10.

3. Rohwer F. Global phage diversity. Cell Press. 2003;113:141.

4. Clokie MRJ, Millard AD, Letarov A V, Heaphy S. Phages in nature. Bacteriophage. 2011;1(1):31–45.

5. Breitbart M, Hewson I, Felts B, Mahaffy JM, Nulton J, Salamon P, et al. Metagenomic Analyses of an Uncultured Viral Community from Human Feces. J Bacteriol. 2003;185(20):6220–3.

6. Manrique P, Bolduc B, Walk ST, Van Der Oost J, De Vos WM, Young MJ. Healthy human gut phageome. Proc Natl Acad Sci. 2016;113(37):10400–5.

7. Shkoporov AN, Hill C. Bacteriophages of the Human Gut: The “Known Unknown” of the Microbiome. Cell Host and Microbe. 2019;Elsevier 25:195–209.

8. Nguyen S, Baker K, Padman BS, Patwa R, Dunstan RA, Weston TA, et al. Bacteriophage transcytosis provides a mechanism to cross epithelial cell layers. MBio. 2017;8(6).

9. Sender R, Fuchs S, Milo R. Are We Really Vastly Outnumbered? Revisiting the Ratio of Bacterial to Host Cells in Humans. Cell. 2016; Cell Press;164:337–40.

10. Barr JJ. A bacteriophages journey through the human body. Immunol Rev. 2017;279(1):106–22.

11. Górski A, Wazna E, Dąbrowska BW, Dąbrowska K, Świtała-Jeleń K, Miȩdzybrodzki R. Bacteriophage translocation. FEMS Immunol Med Microbiol. 2006;46(3):313–9.

12. Ghose C, Ly M, Schwanemann LK, Shin JH, Atab K, Barr JJ, et al. The Virome of Cerebrospinal Fluid: Viruses Where We Once Thought There Were None. Front Microbiol. 2019 Sep;10.

13. Miȩdzybrodzki R, Kłak M, Jonczyk-Matysiak E, Bubak B, Wójcik A, Kaszowska M, et al. Means to facilitate the overcoming of gastric juice barrier by a therapeutic staphylococcal bacteriophage A5/80. Front Microbiol. 2017 Mar;8:1–11.

14. Geier MR, Trigg ME, Merril CR. Fate of bacteriophage lambda in Non-immune germ-1. free mice. Nature. 1973;246(5430):221–3.

15. Huh H, Wong S, St. Jean J, Slavcev R. Bacteriophage interactions with mammalian tissue: Therapeutic applications. Adv Drug Deliv Rev. 2019;145:4–17.

16. Dabrowska K, Switała-Jelen K, Opolski A, Weber-Dabrowska B, Gorski A. A review: Bacteriophage penetration in vertebrates. J Appl Microbiol. 2005;98(1):7–13.

17. Dor-On E, Solomon B. Targeting glioblastoma via intranasal administration of Ff bacteriophages. Front Microbiol. 2015;6.

18. Srivastava AS, Chauhan DP, Carrier E. In utero detection of T7 phage after systemic administration to pregnant mice. BioTechniques. 2004; 37.

19. Gorski A, Dabrowska K, Switala-Jele K, Nowaczyk M, Weber-Dabrowska B, Boratynski J, et al. New insights into the possible role of bacteriophages in host defense and disease. Med Immunol. 2003 Feb;2:2.

20. Karimi M, Mirshekari H, Moosavi Basri SM, Bahrami S, Moghoofei M, Hamblin MR. Bacteriophages and phage-inspired nanocarriers for targeted delivery of therapeutic cargos. Adv Drug Deliv Rev. 2016;106:45–62.

21. Handley SA, Thackray LB, Zhao G, Presti R, Miller AD, Droit L, et al. Pathogenic simian immunodeficiency virus infection is associated with expansion of the enteric virome. Cell. 2012;151(2):253–66.

22. Lehti TA, Pajunen MI, Skog MS, Finne J. Internalization of a polysialic acid-binding Escherichia coli bacteriophage into eukaryotic neuroblastoma cells. Nat Commun. 2017;8(1).

23. Tao P, Mahalingam M, Marasa BS, Zhang Z, Chopra AK, Rao VB. In vitro and in vivo delivery of genes and proteins using the bacteriophage T4 DNA packaging machine. Proc Natl Acad Sci. 2013 Apr;110(15):5846–51.

24. Kerr MC, Teasdale RD. Defining Macropinocytosis. Traffic. 2009 Apr;10(4):364–71.

25. Gordillo Altamirano FL, Barr JJ. Phage Therapy in the Postantibiotic Era. Clin Microbiol Rev. 2019;32(2):1–25.

26. Merril CR, Scholl D, Adhya SL. The prospect for bacteriophage therapy in Western medicine. Nat Rev Drug Discov. 2003 Jun;2(6):489–97.

27. Kutter E, De Vos D, Gvasalia G, Alavidze Z, Gogokhia L, Kuhl S, et al. Phage Therapy in Clinical Practice: Treatment of Human Infections. Curr Pharm Biotechnol. 2010 Feb;11(1):69–86.

28. Abedon ST, Kuhl SJ, Blasdel BG, Kutter EM. Phage treatment of human infections. Bacteriophage. 2011;1(2):66–85.

29. Matsuzaki S, Uchiyama J. Phage Pharmacokinetics: Relationship with Administration Route. In: Phage Therapy: A Practical Approach. Springer International Publishing. 2019;43–57.

30. Dąbrowska K. Phage therapy: What factors shape phage pharmacokinetics and bioavailability? Systematic and critical review. 2019;Med Res Rev 39(5):2000–25.

31. Dąbrowska K, Abedon ST. Pharmacologically Aware Phage Therapy: Pharmacodynamic and Pharmacokinetic Obstacles to Phage Antibacterial Action in Animal and Human Bodies. Microbiol Mol Biol Rev. 2019;83(4):1–25.

32. Payne RJH, Jansen VAA. Understanding bacteriophage therapy as a density-dependent kinetic process. J Theor Biol. 2001 Jan;208(1):37–48.

33. Payne RJH, Jansen VAA. Phage therapy: The peculiar kinetics of self-replicating pharmaceuticals. Clinical Pharmacology and Therapeutics. 2000;68:225–30.

34. Hoffmann M. Animal Experiments on Mucosal Passage and Absorption Viraemia of T3 Phages after Oral, Trachéal and Rectal Administration. Zentralblatt fur Bakteriol Parasitenkunde, Infekt und Hyg. 1965;198(4):371–90.

35. Hildebrand GJ, Wolochow H. Translocation of Bacteriophage Across the Intestinal Wall of the Rat. Exp Biol Med. 1962 Jan;109(1):183–5.

36. Keller R, Engley FB. Fate of Bacteriophage Particles Introduced into Mice by Various Routes. Exp Biol Med. 1958 Jul;98(3):577–80.

37. Sweere JM, Belleghem JD Van, Ishak H, Bach MS, Popescu M, Sunkari V, et al. of bacterial infection. 2019 Mar;9691.

38. Lin Y-W, Kyung Chang RY, Rao GG, Jermain B, Han M-L, Zhao J, et al. Pharmacokinetics/pharmacodynamics of antipseudomonal bacteriophage therapy in rats: A Proof-of-Concept study. Clin Microbiol Infect. 2020 May.

39. Doub JB, Ng VY, Johnson AJ, Slomka M, Fackler J, Horne B, et al. Salvage Bacteriophage Therapy for a Chronic MRSA Prosthetic Joint Infection. Antibiotics. 2020;9:241.

40. Hodyra-Stefaniak K, Lahutta K, Majewska J, Kaźmierczak Z, Lecion D, Harhala M, et al. Bacteriophages engineered to display foreign peptides may become short-circulating phages. Microb Biotechnol. 2019 Jul;12(4):730–41.

41. Carroll-Portillo A, Lin HC. Bacteriophage and the innate immune system: Access and signaling. Microorganisms. 2019;7(12):1–11.

42. Van Belleghem JD, Dąbrowska K, Vaneechoutte M, Barr JJ, Bollyky PL. Interactions between bacteriophage, bacteria, and the mammalian immune system. Viruses. 2019 Jan;11(1).

43. Hodyra-Stefaniak K, Miernikiewicz P, Drapała J, Drab M, Jonczyk-Matysiak E, Lecion D, et al. Mammalian Host-Versus-Phage immune response determines phage fate in vivo. Sci Rep. 2015 Oct;5(1):1–13.

44. Sweere JM, Van Belleghem JD, Ishak H, Bach MS, Popescu M, Sunkari V, et al. Bacteriophage trigger antiviral immunity and prevent clearance of bacterial infection. Science. 2019 Mar;363(6434).

45. Bonilla N, Rojas MI, Netto Flores Cruz G, Hung S-H, Rohwer F, Barr JJ. Phage on tap–a quick and efficient protocol for the preparation of bacteriophage laboratory stocks. PeerJ. 2016;4:e2261.

46. Wang H, Song M. Ckmeans.1d.dp: Optimal k-means clustering in one dimension by dynamic programming. R J. 2011;3(2):29–33.

47. Kim HJ, Li H, Collins JJ, Ingber DE. Contributions of microbiome and mechanical deformation to intestinal bacterial overgrowth and inflammation in a human gut-on-a-chip. Proc Natl Acad Sci. 2016;113(1):e7–15.

48. Kim HJ, Huh D, Hamilton G, Ingber DE. Human Gut-on-a-Chip inhabited by microbial flora that experiences intestinal peristalsis-like motions and flow. R Soc Chem. 2012;12:2165–74.

49. Navabi N, McGuckin MA, Linden SK. Gastrointestinal Cell Lines Form Polarized Epithelia with an Adherent Mucus Layer when Cultured in Semi-Wet Interfaces with Mechanical Stimulation. PLoS One. 2013;8(7).

50. Kim L, Toh YC, Voldman J, Yu H. A practical guide to microfluidic perfusion culture of adherent mammalian cells. Lab Chip. 2007;7(6):681–94.

51. Thuenauer R, Rodriguez-Boulan E, Rümer W. Microfluidic approaches for epithelial cell layer culture and characterisation. Analyst. 2014;139(13):3206–18.

52. Son Y. Determination of shear viscosity and shear rate from pressure drop and flow rate relationship in a rectangular channel. Polymer (Guildf). 2007 Jan;48(2):632–7.

53. Chin WH, Barr JJ. Phage research in ‘organ-on-chip’ devices. Microbiol Aust. 2019 Mar.

54. Barr JJ, Auro R, Sam-Soon N, Kassegne S, Peters G, Bonilla N, et al. Subdiffusive motion of bacteriophage in mucosal surfaces increases the frequency of bacterial encounters. Proc Natl Acad Sci. 2015;112(44):13675–80.

55. Yum K, Hong SG, Healy KE, Lee LP. Physiologically relevant organs on chips. Biotechnol J. 2014 Jan;9(1):16–27.

56. Wang C, Lu H, Alexander Schwartz M. A novel in vitro flow system for changing flow direction on endothelial cells. J Biomech. 2012;45(7):1212–8.

57. Abaci HE, Shen YI, Tan S, Gerecht S. Recapitulating physiological and pathological shear stress and oxygen to model vasculature in health and disease. Sci Rep. 2014 May;4(1):1–9.

58. Park JY, White JB, Walker N, Kuo CH, Cha W, Meyerhoff ME, et al. Responses of endothelial cells to extremely slow flows. Biomicrofluidics. 2011 Jun;5(2):022211.

59. Davies PF, Spaan JA, Krams R. Shear stress biology of the endothelium. Ann Biomed Eng. 2005 Dec;33:1714–8.

60. McQuin C, Goodman A, Chernyshev V, Kamentsky L, Cimini BA, Karhohs KW, et al. CellProfiler 3.0: Next-generation image processing for biology. PLoS Biol. 2018;

61. Hosta-Rigau L, Städler B. Shear stress and its effect on the interaction of myoblast cells with nanosized drug delivery vehicles. Mol Pharm. 2013 Jul;10(7):2707–12.

62. Han J, Zern BJ, Shuvaev V V., Davies PF, Muro S, Muzykantov V. Acute and chronic shear stress differently regulate endothelial internalization of nanocarriers targeted to platelet-endothelial cell adhesion molecule-1. ACS Nano. 2012 Oct;6(10):8824–36.

63. Canton J. Macropinocytosis: New Insights Into Its Underappreciated Role in Innate Immune Cell Surveillance. Front Immunol. 2018;9:2286.

64. Talman L, Agmon E, Peirce SM, Covert MW. Multiscale models of infection. Vol. 11, Current Opinion in Biomedical Engineering. 2019;Elsevier 11:102–8.

65. Bodner K, Melkonian AL, Barth AIM, Kudo T, Tanouchi Y, Covert MW. Engineered Fluorescent E. coli Lysogens Allow Live-Cell Imaging of Functional Prophage Induction Triggered inside Macrophages. Cell Syst. 2020;10(3):254–264.

66. Swanson JA, Watts C. Macropinocytosis. Trends in Cell Biology. 1995;Elsevier Current Trends 5:424–8.

67. Lu F, Wu S-H, Hung Y, Mou C-Y. Size Effect on Cell Uptake in Well-Suspended, Uniform Mesoporous Silica Nanoparticles. Small. 2009; 5(12):1408–13.

68. Yin Win K, Feng SS. Effects of particle size and surface coating on cellular uptake of polymeric nanoparticles for oral delivery of anticancer drugs. Biomaterials. 2005; 26(15):2713–22.

69. Zhu J, Liao L, Zhu L, Zhang P, Guo K, Kong J, et al. Size-dependent cellular uptake efficiency, mechanism, and cytotoxicity of silica nanoparticles toward HeLa cells. Talanta. 2013;107:408–15.

70. Agarwal R, Singh V, Jurney P, Shi L, Sreenivasan S V., Roy K. Mammalian cells preferentially internalize hydrogel nanodiscs over nanorods and use shape-specific uptake mechanisms. Proc Natl Acad Sci. 2013 Oct;110(43):17247–52.

71. Hsiao I-L, Gramatke AM, Joksimovic R, Sokolowski M, Gradzielski M, Haase A. Size and Cell Type Dependent Uptake of Silica Nanoparticles. J Nanomed Nanotechnol. 2014;05(06).

72. Kellermayer MSZ, Vörös Z, Csík G, Herényi L. Forced phage uncorking: viral DNA ejection triggered by a mechanically sensitive switch. Nanoscale. 2018;10:1898.

73. Barr JJ, Auro R, Furlan M, Whiteson KL, Erb ML, Pogliano J, et al. Bacteriophage adhering to mucus provide a non-host-derived immunity. Proc Natl Acad Sci. 2013;110(26):10771–6.

74. Geier MR, Trigg ME, Merril CR. Fate of bacteriophage lambda in Non-immune germ-free mice. Nature. 1973;246(5430):221–3.

75. Merril CR, Biswas B, Carlton R, Jensen NC, Creed GJ, Zullo S, et al. Long-circulating bacteriophage as antibacterial agents. Proc Natl Acad Sci. 1996 Apr;93(8):3188–92.

76. Schooley RT, Biswas B, Gill JJ, Hernandez-Morales A, Lancaster J, Lessor L, et al. Development and use of personalized bacteriophage-based therapeutic cocktails to treat a patient with a disseminated resistant Acinetobacter baumannii infection. Antimicrob Agents Chemother. 2017;61(10):1–14.

77. Schindelin J, Arganda-Carreras I, Frise E, Kaynig V, Longair M, Pietzsch T, et al. Fiji: an open-source platform for biological-image analysis. Nat Methods. 2012;9(7).

78. Bunn A, Korpela M. A dendrochronology program library in R (dplR). Dendrochronologia. 2018 Jul;26(2):115–24.

