## Supplemental Data 2_RServer for "Bacteriophage uptake by Eukaryotic cell layers represents a major sink for phages during therapy": SD2_server.R

```
library(mrgsolve)
library(ggplot2)
library(dplyr)
library(shiny)

shinyServer(function(input, output) {
   
  SData <- reactive({
    
    code <- '
    $PARAM
    TVVMAX = 1, TVVC=1, TVVP1 = 1, TVVP2 = 1, TVQ1= 1, TVQ2 = 1, TVKM = 1, TVKD1 = 1, TVKD2 = 1
    
    $CMT CENT P1 P2 R
    
    $MAIN
    double VMAX  = TVVMAX  * exp(ETA(1));
    double VC    = TVVC    * exp(ETA(2));
    double VP1   = TVVP1   * exp(ETA(3));
    double VP2   = TVVP2   * exp(ETA(4));
    double Q1    = TVQ1    * exp(ETA(5));
    double Q2    = TVQ2    * exp(ETA(6));
    double KM    = TVKM    * exp(ETA(7));
    
    double KD1   = TVKD1;
    double KD2   = TVKD2;
 
    $OMEGA  
    0 0 0 0 0 0 0
    
    $ODE
    
    dxdt_CENT = - Q1*(CENT/VC) + Q1*(P1/VP1) - Q2*(CENT/VC) + Q2*(P2/VP2) - KD1*CENT + KD2*R - VMAX*(CENT/VC) /((CENT/VC) + KM) ;
    dxdt_P1   =   Q1*(CENT/VC) - Q1*(P1/VP1);
    dxdt_P2   =                                Q2*(CENT/VC) - Q2*(P2/VP2);
    dxdt_R    =                                                             KD1*CENT - KD2*R;
    
    $TABLE
    
    double IPRED = CENT/VC;
    double DV = log10(IPRED + 0.0001);
    
    $CAPTURE DV
   
  '
    
    #Load model
    mod <- mcode_cache("PK", code)
    
    ## simulation settings
    nsamples <- input$nsamples
    simtime <- input$simtime
    delt <-input$delta
    
    #Dose settings
    Dose     <- input$DOSE
    DTime    <- input$DTIME
    Interval <- input$INTERVAL
    Addl     <- input$ADDL
    
    ## THETA  
    VMAX  <- input$VMAX 
    VC    <- input$VC 
    VP1   <- input$VP1
    VP2   <- input$VP2
    Q1    <- input$Q1
    Q2    <- input$Q2
    KM    <- input$KM
    KD1   <- input$KD1
    KD2   <- input$KD2
        
    #INTER-INDIVIDUAL VARIABILITY (ETA)
    ETAVMAX   <- input$ETAVMAX 
    ETAVC     <- input$ETAVC
    ETAVP1    <- input$ETAVP1 
    ETAVP2    <- input$ETAVP2
    ETAQ1     <- input$ETAQ1 
    ETAQ2     <- input$ETAQ2
    ETAKM     <- input$ETAKM 
    
    ## RANDOM UNEXPLAINED VARIABILITY (SIGMA)
    ADD  <- input$ADD

    ## Dosing Objects + Create dataset for simulations
    obj <- as.data.frame(ev(ID = 1:nsamples, ii = Interval, rate = 0, addl = Addl, amt = Dose, time = DTime))
    dat <- obj[order(obj$ID, obj$time),]
    dat$TVVMAX  <- VMAX
    dat$TVVC    <- VC
    dat$TVVP1   <- VP1
    dat$TVVP2   <- VP2
    dat$TVQ1    <- Q1
    dat$TVQ2    <- Q2
    dat$TVKM    <- KM
    dat$TVKD1     <- KD1
    dat$TVKD2     <- KD2

    ## Omege Matrix
    omega <- dmat(ETAVMAX,ETAVC,ETAVP1,ETAVP2,ETAQ1,ETAQ2,ETAKM)
    
    ## Simga matrix 
    #sigma <- dmat(ADD)
    
    res <- mod %>%
      omat(omega) %>%
      data_set(dat) %>%
      mrgsim(end= simtime, obsonly=TRUE, delta=delt) 
    
    outputdata <- as.data.frame(res)
  
    
    return(outputdata)
    
  })
  
  cal <- reactive({
    
    cal <- SData() %>%
      group_by(time) %>%
      summarise(Out=median(DV)) 
    
    return(cal)
    
  })

  
  output$Plot <- renderPlot({
    
    ggplot(cal(), aes(x=time,y=Out)) +
      
      scale_x_continuous(expand = c(0, 0)) + scale_y_continuous(expand = c(0, 0), limits = c(0, 12)) +
      
      geom_line(colour="red") +
      
      xlab("Time(h)") +
      
      ylab("Phage Titer (Log PFU/mL)")+
      
      theme_bw(14) +
      
      theme(legend.position="none")
  })
  
  # Downloadable file
  
  output$downloadData <- downloadHandler(
    filename = function() {
      "simuated_dataset.csv"
    },
    content = function(file) {
      write.csv(SData(), file, row.names = TRUE)
    }
  )
  
  
})

  
  
```
