## Supplemental Data 3_Rui for "Bacteriophage uptake by Eukaryotic cell layers represents a major sink for phages during therapy": SD3_ui.R

```
#=Header-(start)===========================================================
#-------------------------------------------------------------------------+
# Author            : Yu-Wei Lin 
# Creation date     : 01 August 2020 
# Purpose           : <Brief description of the program goal(s)>
#-------------------------------------------------------------------------+
# Platform          : x86_64-w64-mingw32 running Windows >= 8 x64 (build 9200)
# Environment       : R version 3.6.1 (2019-07-05)
#-------------------------------------------------------------------------+

library(shiny)
library(shinythemes)
library(shinydashboard)

shinyUI(fluidPage(theme = shinytheme("superhero"),
                  
header <- dashboardHeader(),

titlePanel("Hypothetical Rat Phage PK Model with In Vitro Inactivation Rate Constant"),
                  
sidebarLayout(position ="right",
sidebarPanel(
h4("Dosage Regimens"),
numericInput("DOSE", label='Dose (PFU/mL)', value = 1000000000, min=0, max=10000000000),

br(),
h4("Phage Inactivation Constant (Hypothetical)"),
numericInput("KD1",  label = 'First-Order Inactivation Constant (1/h/rat)', value = 0,  min=0.00001, max=100),
numericInput("KD2",  label = 'First-Order Re-activation Constant (1/h/rat)', value = 0, min=0.00001, max=100),

br(),
h4("Simulation Settings"),
numericInput("DTIME",    label = 'Dosing Time', value = 0, min = 0),
numericInput("INTERVAL", label = 'Dosing Interval', value = 0, min = 0),
numericInput("ADDL",     label = 'Dosing Frequency', value = 0, min = 0),
numericInput("nsamples", label = 'Number of rats per simulation', min=0, max=2000, value = 1),
numericInput("simtime",  label = 'Simulation time', value = 24,min=0, max = 48),
sliderInput("delta",     label = 'Integration step', value = 0.1, min = 0.001, max=1),

br(),
h4("Theta"),
numericInput("VC",    label = 'Central Volume of Distribution (mL/Rat)', value = 111,min=0, max=100),
numericInput("VMAX",  label = 'Maximum Eliminataion Rate (PFU/h/Rat)', value = 43900000000, min=0, max=100000000),
numericInput("VP1",   label = 'Peripheral Volume of Distribution 1 (mL/Rat)', value = 128,min=0, max=100),
numericInput("VP2",   label = 'Peripheral Volume of Distribution 2 (mL/Rat)', value = 180, min=0, max=100),
numericInput("Q1",    label = 'Intercompartmental Clearance 1 (mL/h/Rat)', value = 30.4,min=0, max=100),
numericInput("Q2",    label = 'Intercompartmental Clearance 2 (mL/h/Rat)', value = 538, min=0, max=1000),
numericInput("KM",    label = '50% of the Maximal Elimination Rate (PFU/mL/Rat)', value = 16400000, min = 0, max=100),

br(),
h4("Inter-animal Variability"),
sliderInput("ETAVC",   label = 'Central Volume of Distribution', value = 0, min=0, max=1),
sliderInput("ETAVMAX", label = 'Maximum Eliminataion Rate', value = 0, min=0, max=1),
sliderInput("ETAVP1",  label = 'Peripheral Volume of Distribution 1', value = 0, min=0, max=1),
sliderInput("ETAVP2",  label = 'Peripheral Volume of Distribution 2', value = 0, min=0, max=1),
sliderInput("ETAQ1",   label = 'Intercompartmental Clearance 1', value = 0, min=0, max=1),
sliderInput("ETAQ2",   label = 'Intercompartmental Clearance 2', value = 0, min=0, max=1),
sliderInput("ETAKM",   label = '50% of the Maximal Elimination Rate', value = 0, min=0, max=1),
            
),
                    
mainPanel(
plotOutput("Plot"),
downloadButton("downloadData", "Export")
   )
      )
          )
)
```
