## Supplemental Figures for "Bacteriophage uptake by Eukaryotic cell layers represents a major sink for phages during therapy"

**Movies from SM1 - SM15 method:**

Phages were applied to the cell layer and incubated for two hours (or one hour for SM8 and SM9 movies) on a μ-Slide 8 well glass bottom slide on a microscope stage with temperature and $CO_2$ control. One image was acquired every two minutes on an inverted Leica SP8 confocal microscope with HC PL APO 63x/1.40 CS2 oil immersion objective. A hybrid detector (HyD) was used in sequential mode to visualise phage DNA. Cells were stained with nucleus stain, Hoescht 33342 (blue), plasma membrane stain, CellMask (magenta) and T4 phages labelled with DNA-complexing stain, SYBR-Gold (green). Scale bar: 10 μm; Timing: hours:minutes.

**SM1. T4-A549.** T4 phage was applied to human epithelial lung cells, A549.

**SM2. T4-HUVEC.** T4 phage was applied to human endothelial cells from the umbilical vein, HUVEC-EC cells.

**SM3. T4-MDCK-I.** T4 phage was applied to fibroblast dog kidney cells, MDCK-I.

**SM4. T4-HeLa.** T4 phage was applied to human cervix epithelial cells, HeLa.

**SM5. T4-THP-1.** T4 phage was applied to human macrophage induced monocytes, THP-1.

**SM6. T4-HT29.** T4 phage was applied to human colon epithelial cells, HT29.

**SM7. T4-BJ.** T4 phage was applied to human skin fibroblast cells, BJ.

**SM8. T4-MDCK-I-3D-Timelapse.** T4 phage was applied to fibroblast dog kidney cells, MDCK-I.

**SM9. T4-MDCK-I-3D.** T4 phage was applied to fibroblast dog kidney cells, MDCK-I, using the Imaris software we 3D reconstituted the volume of the cell. Scale bar: 10 μm.

**SM10. Lambda-A549.** Lambda phage was applied to human epithelial lung cells, A549.

**SM11. Lambda-HUVEC.** Lambda phage was applied to human endothelial umbilical vein cells, HUVEC-EC.

**SM12. Lambda-BJ.** Lambda was applied to human fibroblast skin cells, BJ.

**SM13. T3-A549.** T3 phage was applied to human epithelial lung cells, A549.

**SM14. T3-HUVEC.** T3 phage was applied to human endothelial umbilical vein cells, HUVEC-EC.

**SM15. T3-BJ.** T3 phage was applied to human fibroblast skin, BJ.

**SD1. CellProfiler Pipeline.** Pipeline used in CellProfiler to analyse the greyvalues as a proxy for fluorescence intensity of the phage channel in supplemental figure two and figure 4B.

**SD2_server.R.** R code for the mathematical model.

**SD3_ui.R.** The R code for the user interface of the mathematical model.

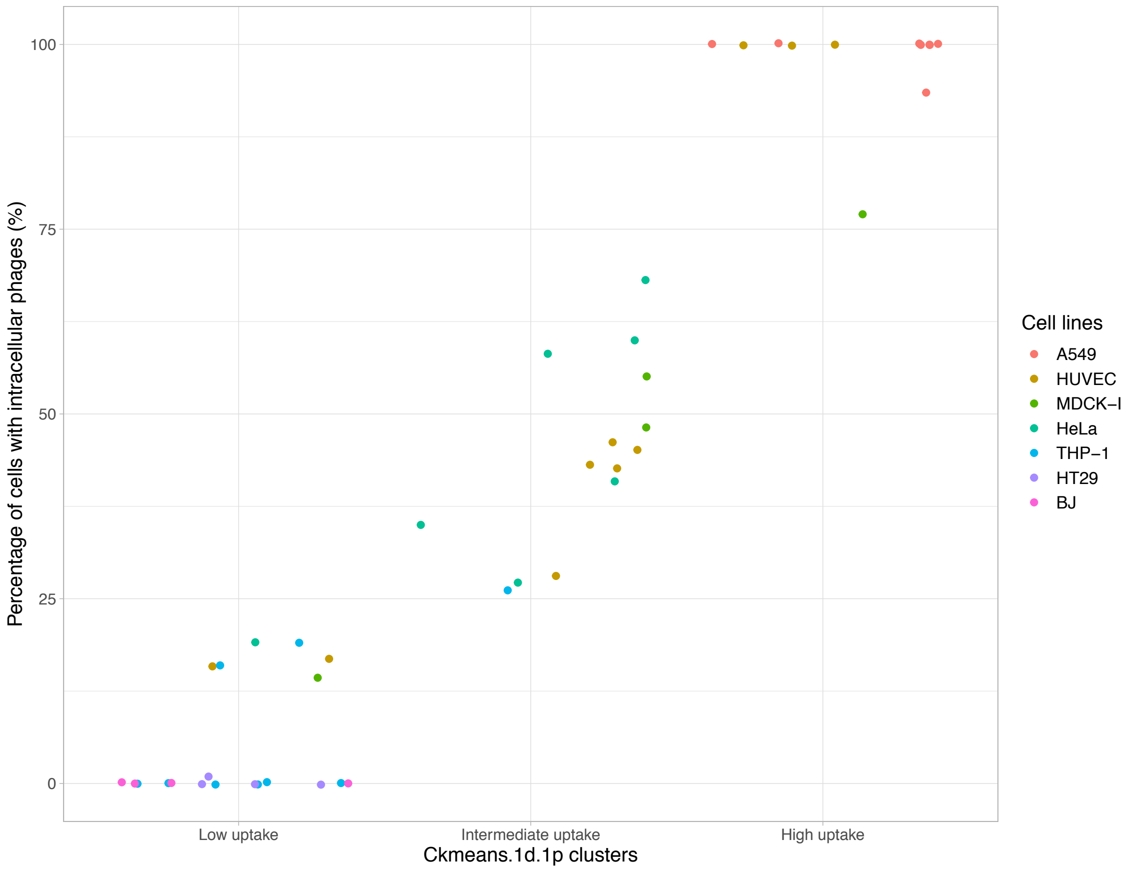


**Supplemental Figure 1.** Clustering was performed using the dynamic programming algorithm in the R package Ckmeans.1d.dp. The given number of clusters was three shown here in the X axis (1 – low uptake, 2 – intermediate uptake and 3 – high uptake) and the percentage of uptake in Y axis. Each colour represents one of the cell lines tested.

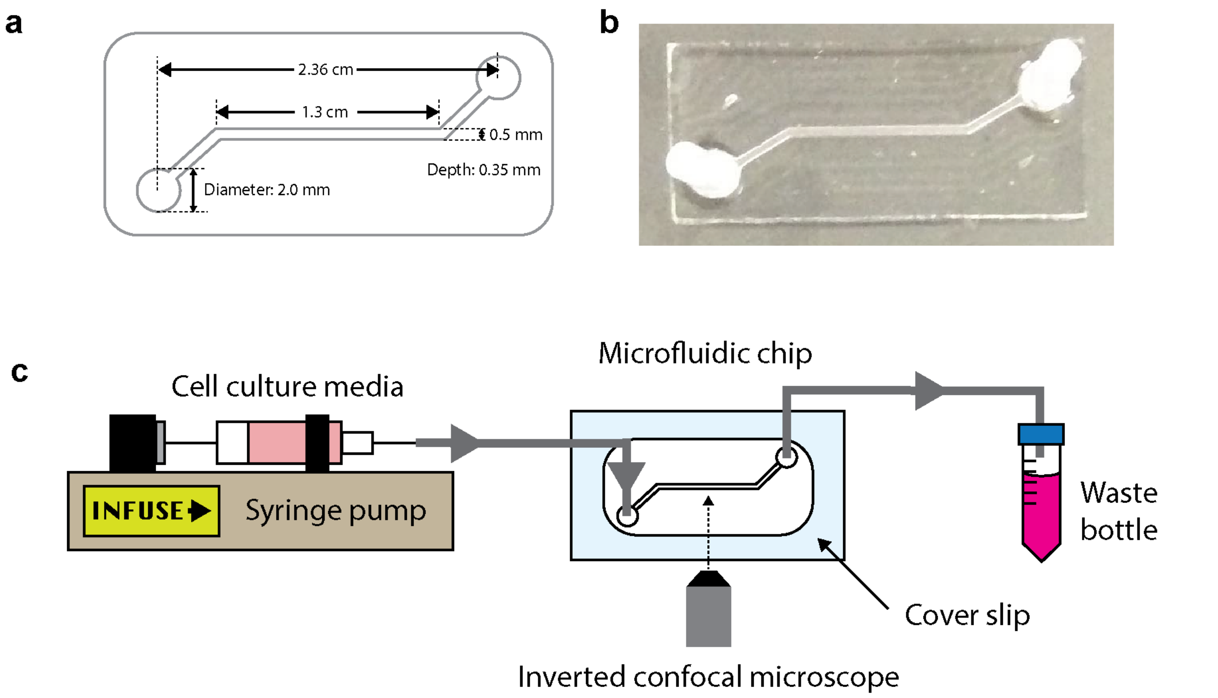


**Supplemental Figure 2**. Microfluidic device and set-up for phage transcytosis experiment in flow conditions. **(a)** Chip channel dimensions, **(b)** the actual device picture. The chip was fabricated using PDMS and irreversibly bonded onto a glass cover slip using plasma. **(c)** Experimental set-up schematic for transcytosis experiment on flow. After inoculating the device with phages, the cell layer within the microfluidic chip was maintained under flow with egressing fluid collected as waste. Phage and cell layer behaviour were monitored and recorded via an inverted confocal microscope.

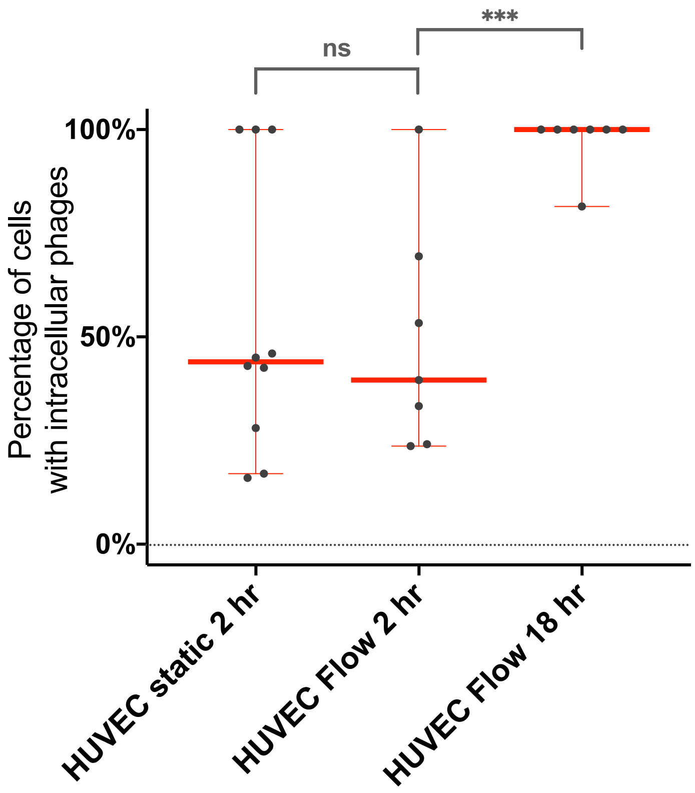


**Supplemental Figure 3**. Percentage of HUVEC cells containing intracellular phages at the two-hour time point under static conditions and the two time points under flow condition of 0.1 dyne/cm$^2$, two hours and 18 hours. Scatter plots show medians of percentage of cells with intracellular phages; error bars represent 95% confidence intervals; each dot represents one Field of View (FOV). (static, *n* = 10; Flow 2 hr, *n* = 7; Flow 18 hours, *n* = 7). P-values calculated from a one-way unpaired t-test (P < 0.001: ***; ns: non-significant).

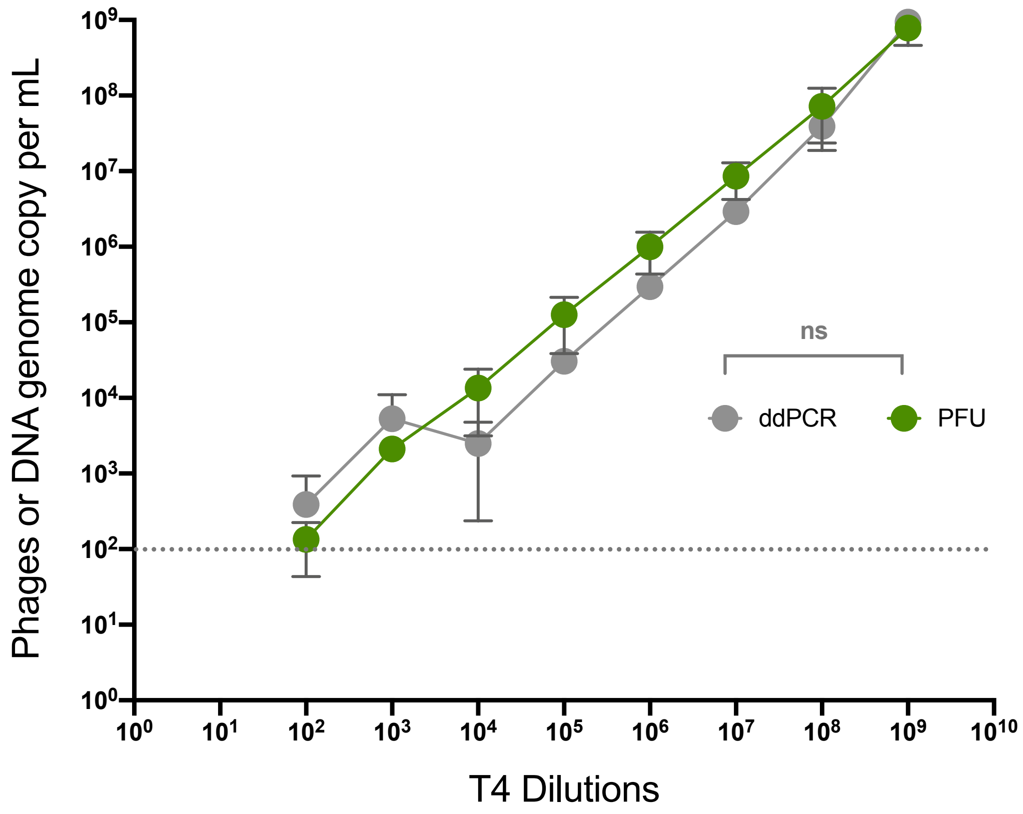


**Supplemental Figure 4**. Dilution series of T4 phage stock using PFU and ddPCR techniques. X axis representing the dilution of the phage stock, in the Y axis the concentrations obtained with PFU or ddPCR in Phages or DNA genome copy per ml respectively. Shown in green line the PFU results and in grey the ddPCR results. Limit of detection is represented by the grey dotted line. P values calculated from paired T-test (ns: non-significant).

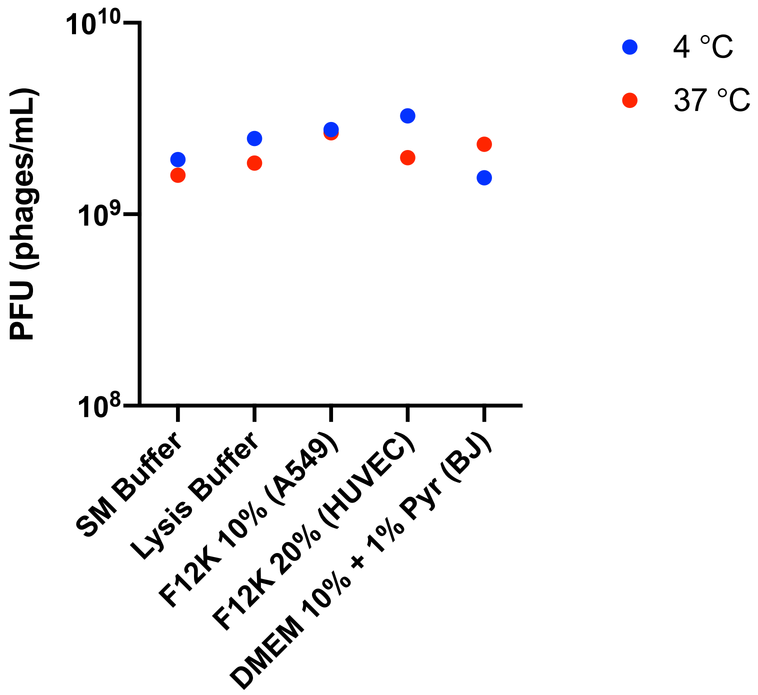


**Supplemental Figure 5**. Concentrations of T4 phage stock using PFU techniques. X axis representing the different media of incubation: stocking media (SM Buffer), the lysis buffer used to lyse the cells and collect the intracellular phages, and the three spent cell culture media used to grow A549, HUVEC or BJ cells. The spent cell culture media was collected after two days of cells growth before being used in this control. In the Y axis the concentrations obtained with PFU in phages/ml. Shown in blue dots the results at 4 °C and in red dots the results at 37 °C. Non-significant P values calculated from one-way ANOVA.

**Supplemental Table 1.** Shear stress calculation.

| | Constant | μ: Viscosity of media at 37 °C (mPa·s) | Q: flow rate ($m^3/s$) | W: channel width (m) | H: channel height (m) | T: sheer stress ($dyne/cm^2$) |
|---|---|---|---|---|---|---|
| Values | 6 | 0.78 | $1.33 \times 10^{-10}$ | $5 \times 10^{-4}$ | $3.5 \times 10^{-4}$ | $T = \frac{6\mu Q}{WH^2}$ |

**Supplemental Table 2.** Primers and probe DNA sequences for gp23 ddPCR assay.

| **Primers/probes** | **Sequences** | **Vendor** |
|---|---|---|
| Primer gp23 Forward | 5'-CTGCAGGTCAGACTTCTG-3' | Micromon |
| Primer gp23 Reverse | 5'-CATCGGCTGAACACCAC-3' | Micromon |
| Probe gp23 | 5'-**56-FAM**/ACTCAGATT/**ZEN**/GGCCCAGCTGTT/**3IABkFQ**/-3' | Integrated DNA Technology (IDT) |

**Supplemental Table 3.** PCR mix for gp23 ddPCR assay.

| | **For one reaction (µl)** | **For 8 reactions (µl)** |
|---|---|---|
| Super Mix for Probes | 10 | 80 |
| Forward gp23 Primer 10 µM | 1.8 | 14.4 |
| Reverse gp23 Primer 10 µM | 1.8 | 14.4 |
| Gp23 probe 20X | 1 | 8 |
| Water | 5.4 | 43.2 |
| Sample | 2 | - |
| Total | 22 | 20 mix + 2 µl sample |

**Supplemental Table 4.** Thermal cycling conditions for gp23 ddPCR assay.

| **Cycling steps** | **Time (hr:min:sec)** | **Temperatures (°C)** | **Number of cycles** |
|---|---|---|---|
| Enzyme activation | 00:10:00 | 95 | 1 |
| Denaturation | 00:00:30 | 94 | 40 |

| | | | |
|---|---|---|---|
| Annealing | 00:01:30 | 55 | 40 |
| Extension | 00:00:30 | 72 | 40 |
| Enzyme deactivation | 00:10:00 | 98 | 1 |
| Hold | ∞ | 4 | 1 |

**Supplemental Tables 5.** Calculation for math model and constants used.

Calculation of the first order constant:

$$\frac{d[ddPCR]}{dt} = k[ddPCR]$$

$$k = \frac{\ln[ddPCR_o] - \ln[ddPCR]}{t}$$

| | |
|---|---|
| $[ddPCR]$ | $phage\ conc.\ in\ central\ compartment$ |
| $[ddPCR_o]$ | $initial\ phage\ conc.\ administered$ |
| $t$ | $elapsed\ time$ |
| $k$ | $rate\ constant$ |
| $[ddPCR_o]$ *at t:0 (DNA genome copies/mL)* | *1000000000* |
| $[ddPCR]$ *at t:30 seconds (DNA genome copies/mL) ** | *3392* |
| $[ddPCR]$ *at t:18 hours (DNA genome copies/mL) ** | *363073* |

*Geometric mean of ddPCR data combined between each replicates and cells lines

Constants used in the model:

| Data from | Lin et al. 2020 |
|---|---|
| Central volume of distribution (mL/rat) | 111 |
| Maximum Elimination rate (PFU/h/rat) | 43900000000 |
| Peripheral volume of distribution 1 (mL/rat) | 128 |
| Peripheral volume of distribution 2 (mL/rat) | 180 |
| Intercompartmental clearance 1 (mL/h/rat) | 30.4 |
| Intercompartmental clearance 2 (mL/h/rat) | 538 |

| | |
|---|---|
| 50% of the maximal elimination rate (PFU/mL/rat) | 16400000 |
| Dosing time | 1 |
| Number of rats per simulation | 1 |
| Simulation time (h) | 50 |
| Integration step | 0.001 |
| Phage initial dose (PFU/mL) | 1000000000 |
| First-order inactivation constant t:0 (1/h/rat) | 0 |
| First-order inactivation constant t:30 seconds (1/h/rat) | 1511 |
| First-order inactivation constant t:18 hours (1/h/rat) | 0.44 |
| Formula used to plot the graph in Fig. 6B | DV = log10(CENT/VC + 0.0001) |
